# Radiogenomics of *C9orf72* expansion carriers reveals global transposable element de-repression and enables prediction of thalamic atrophy and clinical impairment

**DOI:** 10.1101/2022.07.28.501897

**Authors:** Luke W. Bonham, Ethan G. Geier, Daniel W. Sirkis, Josiah K. Leong, Eliana Marisa Ramos, Qing Wang, Anna Karydas, Suzee E. Lee, Virginia E. Sturm, Russell P. Sawyer, Adit Friedberg, Justin K. Ichida, Aaron D. Gitler, Leo Sugrue, Michael Cordingley, Walter Bee, Eckard Weber, Joel Kramer, Katherine P. Rankin, Howard J. Rosen, Adam L. Boxer, William W. Seeley, John Ravits, Bruce L. Miller, Jennifer S. Yokoyama

## Abstract

Hexanucleotide repeat expansion (HRE) within *C9orf72* is the most common genetic cause of frontotemporal dementia (FTD). Thalamic atrophy occurs in both sporadic and familial FTD but is thought to distinctly affect HRE carriers. Separately, emerging evidence suggests widespread de-repression of transposable elements (TEs) in the brain in several neurodegenerative diseases, including *C9orf72* HRE-mediated FTD (C9-FTD). Whether TE activation can be measured in peripheral blood and how the reduction in peripheral *C9orf72* expression observed in HRE carriers relates to atrophy and clinical impairment remain unknown. We used the FreeSurfer pipeline and its extensions to assess the effects of *C9orf72* HRE and clinical diagnosis (*n* = 78) on atrophy of thalamic nuclei. We also generated a novel, whole-blood RNA-seq dataset to determine the relationships between peripheral *C9orf72* expression, TE activation, thalamic atrophy, and clinical severity (*n* = 114). We confirmed global thalamic atrophy and reduced *C9orf72* expression in HRE carriers. Moreover, we identified disproportionate atrophy of the right mediodorsal lateral nucleus in HRE carriers and showed that *C9orf72* expression associated with clinical severity, independent of thalamic atrophy. Strikingly, we found global peripheral activation of TEs, including the human endogenous LINE-1 element, *L1HS. L1HS* levels were associated with atrophy of multiple pulvinar nuclei, a thalamic region implicated in C9-FTD. Integration of peripheral transcriptomic and neuroimaging data from HRE carriers revealed atrophy of specific thalamic nuclei; demonstrated that *C9orf72* levels relate to clinical severity; and identified marked de-repression of TEs, including *L1HS*, which predicted atrophy of FTD-relevant thalamic nuclei.

**Significance Statement:** Pathogenic repeat expansion in *C9orf72* is the most frequent genetic cause of frontotemporal dementia and amyotrophic lateral sclerosis (C9-FTD/ALS). The clinical, neuroimaging, and pathological features of C9-FTD/ALS are well-characterized, whereas the intersections of transcriptomic dysregulation and brain structure remain largely unexplored. Herein, we utilized a novel radiogenomic approach to examine the relationship between peripheral blood transcriptomics and thalamic atrophy, a neuroimaging feature disproportionately impacted in C9-FTD/ALS. We confirmed reduction of *C9orf72* in blood and found broad dysregulation of transposable elements—genetic elements typically repressed in the human genome—in symptomatic *C9orf72* expansion carriers, which associated with atrophy of thalamic nuclei relevant to FTD. *C9orf72* expression was also associated with clinical severity, suggesting that peripheral *C9orf72* levels capture disease-relevant information.

## Introduction

Hexanucleotide repeat expansion (HRE) intronic to *C9orf72* is the most common genetic cause of frontotemporal dementia (FTD) and amyotrophic lateral sclerosis (ALS). Since the association of *C9orf72* HRE with FTD and ALS over a decade ago (Dejesus-Hernandez et al., 2011; Renton et al., 2011), several classes of pathogenic mechanisms have been characterized and invoked to explain HRE pathogenicity. These mechanisms are categorized broadly as involving gains of toxic function and partial loss of C9orf72 protein function (reviewed in (Balendra and Isaacs, 2018; Vatsavayai et al., 2018; Braems et al., 2020)). Gain-of-function (GOF) mechanisms include the formation of repeat-containing RNA foci thought to result in sequestration of RNA-binding proteins, and the generation of dipeptide repeat proteins non-canonically translated from the expanded GGGGCC (G_4_C_2_) repeat present within the *C9orf72* mRNA of HRE carriers. Haploinsufficiency, on the other hand, has received increasing interest in the last several years and has been proposed to act synergistically with GOF mechanisms to drive pathogenicity in expansion carriers. Precisely how these putative mechanisms lead to loss of nuclear TAR DNA-binding protein 43 (TDP-43) and TDP-43 aggregation that is characteristic of *C9orf72*-associated ALS/FTD neuropathology remains unknown.

Neuroanatomically, the earliest cortical regions thought to be affected in the behavioral variant of FTD (bvFTD), which impacts social behavior and emotional processing, are anterior cingulate and frontoinsular cortices (Seeley et al., 2008, 2012). However, prominent atrophy of the thalamus also occurs in both sporadic and familial forms of FTD (Bocchetta et al., 2018), including FTD due to *C9orf72* HRE (hereafter, C9-FTD) as well as pathogenic variation in *MAPT, GRN*, and other FTD-associated genes. Intriguingly, several lines of evidence suggest that the thalamus may be disproportionately affected in C9-FTD, and specific nuclei such as the medial pulvinar may be uniquely affected (Sha et al., 2012; Yokoyama and Rosen, 2012; Lee et al., 2014; Yokoyama et al., 2014; Vatsavayai et al., 2016; Bocchetta et al., 2020).

The discovery that repetitive element (RE) transcripts are elevated in C9-ALS/FTD brains (Prudencio et al., 2017) and that loss of nuclear TDP-43 is associated with de-condensation of REs such as long interspersed nuclear elements (LINEs) and consequent LINE1 retrotransposition (Liu et al., 2019) has provided an additional pathobiological mechanism to consider. This phenomenon is unlikely to be restricted to C9-ALS/FTD—a subset of postmortem brain tissue from sporadic ALS also exhibits a profile of increased retrotransposon expression reminiscent of that observed in C9-ALS/FTD (Tam et al., 2019). Moreover, transposable element (TE) activation may occur in the context of other proteinopathies as well—tau neuropathology also appears to induce TE expression (Guo et al., 2018; Sun et al., 2018). Remarkably, a *Drosophila* model of pathogenic *CHMP2B* variation, which causes FTD characterized by atypical TDP-43- and tau-negative neuropathology (Skibinski et al., 2005; Mackenzie and Neumann, 2016), also involves augmented TE expression (Fort-Aznar et al., 2020), suggesting that heightened TE expression and retrotransposition may represent a general mechanism underlying multiple forms of neurodegeneration. However, to our knowledge, it is unknown whether this activation occurs in the periphery outside of the brain.

In this paper, we integrated peripheral blood RNA sequencing (RNA-seq) data and neuroimaging analyses from *C9orf72* HRE carriers from across the clinical spectrum and healthy controls to (i) confirm global thalamic atrophy and reduced peripheral expression of *C9orf72* in HRE carriers; (ii) identify disproportionate atrophy of specific thalamic nuclei in HRE carriers; (iii) show that peripheral *C9orf72* expression associates with clinical impairment independent of thalamic atrophy; (iv) discover global peripheral de-repression of TEs in affected HRE carriers; (v) demonstrate strikingly increased expression of the human-specific LINE1 element, *L1HS*, in symptomatic HRE carriers; and (vi) show that peripheral *L1HS* levels associate with thalamic nuclei volumes in FTD-relevant regions. Our results indicate that de-repression of TE expression in C9-ALS/FTD patients is not restricted to the central nervous system (CNS). Peripheral upregulation of TEs such as *L1HS* may therefore enable novel, blood-based biomarkers for C9-ALS/FTD.

## Materials and Methods

### Study Participants

All participants or their surrogates provided written informed consent prior to study participation, and all aspects of the studies described here were approved by the UCSF or UCSD institutional review boards (IRBs).

#### (i) Neuroimaging study

Seventy-eight individuals (*n* = 44 cognitively normal controls and *n* = 34 *C9orf72* HRE carriers) participated in this study. Individuals were recruited from the San Francisco Bay Area as part of ongoing studies of normal aging and FTD at the University of California, San Francisco (UCSF) Memory and Aging Center (MAC). *C9orf72* HRE carriers had a range of clinical diagnoses seen across the spectrum of *C9orf72* HRE-related disease, including cognitively normal (presymptomatic; *n* = 10), mild cognitive impairment (MCI; *n* = 7), bvFTD (*n* = 13), and bvFTD with motor neuron disease (*n* = 4). There were significant differences between normal controls and *C9orf72* HRE carriers when compared by age and education (see Table 1).

**Table 1.**
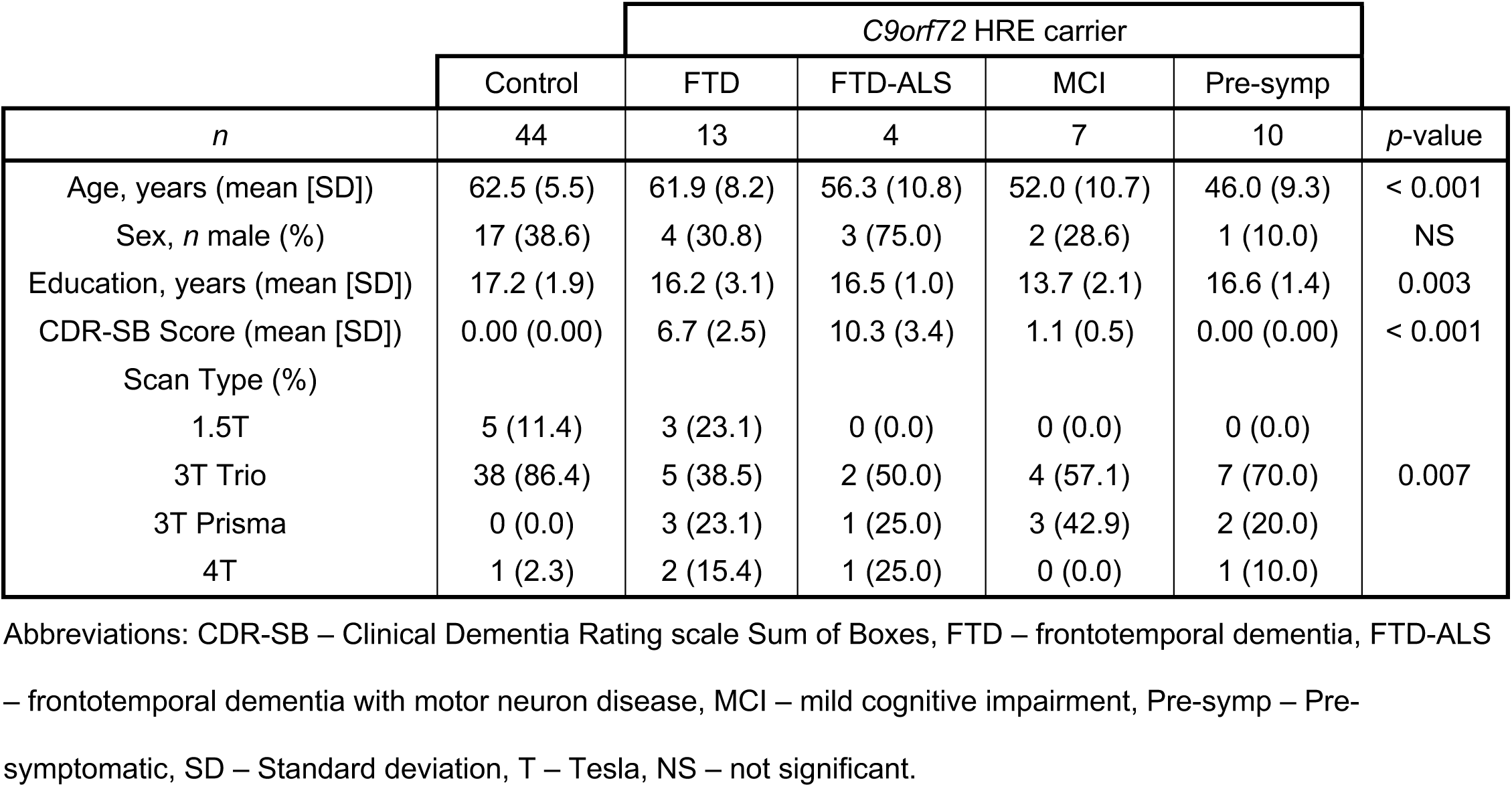
Demographic characteristics of neuroimaging cohort.

#### (ii) RNA-seq study

For whole-blood donors, participants carrying a pathogenic HRE in *C9orf72*, defined as more than 30 repeats (*n* = 49; *C9orf72*+) (Renton et al., 2011), were assessed and clinically diagnosed at the UCSF MAC. No participants in this study carried other known neurodegenerative disease-causing pathogenic variants. Participants with mild cognitive or behavioral symptoms were classified as having MCI, while *C9orf72*+ participants who did not display any symptoms were classified as pre-symptomatic. Cognitively normal, healthy older adult controls (*n* = 65; mean age = 61.3 ± 6.7 years) were recruited to the UCSF MAC as part of ongoing longitudinal studies of aging. Demographic information for participants in this study is included in Table 2. Differences in participant demographics were assessed by one-way analysis of variance (ANOVA) followed by Tukey’s test for post-hoc analysis, or chi-square test. A *p* < 0.05 was considered statistically significant. For the PBMC RNA-seq study, patients diagnosed with ALS met the modified El Escorial criteria for ALS (Brooks et al., 2000) (see Extended Data Table 2-1 for demographic information).

**Table 2.** Demographic characteristics of RNA-seq cohort.

|  |  | *C9orf72* HRE carrier |  |  |  |  |
| --- | --- | --- | --- | --- | --- | --- |
|  | Control | FTD | FTD-ALS | MCI | Pre-symp |  |
| *n* | 65 | 20 | 7 | 10 | 12 | *p*-value |
| Age, years (mean [SD]) | 61.3 (6.7) | 60.6 (8.2) | 57.7 (11.2) | 57.5 (13.1) | 45.7 (9.2) | $< 1 \times 10^{-5}$ |
| Sex, *n* male (%) | 27 (41.5) | 6 (30.0) | 5 (71.4) | 5 (50.0) | 3 (25.0) | NS |
| CDR-SB Score (mean [SD]) | 0 (0) | 7.6 (2.9) | 8.4 (3.5) | 1.3 (0.9) | 0 (0) | $< 1 \times 10^{-5}$ |

Abbreviations: CDR-SB – Clinical Dementia Rating scale Sum of Boxes, FTD – frontotemporal dementia, FTD-ALS – frontotemporal dementia with motor neuron disease, MCI – mild cognitive impairment, Pre-symp – Pre-symptomatic, SD – Standard deviation, NS – not significant.

### Clinical Assessment

Study participants underwent multistep screening prior to an in-person clinical evaluation at the UCSF MAC that included a neurological exam, cognitive assessment, and medical history (Rankin et al., 2005; Miller et al., 2013). In addition, each participant’s study partner was interviewed regarding the participant’s functional abilities. A multidisciplinary team composed of a behavioral neurologist, neuropsychologist, and registered nurse then established clinical diagnoses for cases according to consensus criteria for MCI (Petersen et al., 1999, 2014), FTD and its subtypes (Gorno-Tempini et al., 2011; Rascovsky et al., 2011), ALS (Brooks et al., 2000), and FTD-ALS (Strong et al., 2017). Controls and pre-symptomatic *C9orf72* HRE carriers had a Clinical Dementia Rating scale Sum of Boxes (CDR-SB) (Morris, 1993) score of 0, a Mini Mental State Exam (MMSE) (Folstein et al., 1975) score > 26, and a normal neurological exam.

### Neuroimaging

Study participants underwent structural T1-weighted magnetic resonance imaging (MRI) at one of two imaging centers at UCSF: 65 participants were scanned at the UCSF Neuroscience Imaging Center (NIC) (*n* = 38 controls, *n* = 27 *C9orf72* HRE carriers) on a 3T scanner while 13 participants (*n* = 7 controls, *n* = 6 *C9orf72* HRE carriers) were imaged at the San Francisco Veterans Affairs Medical Center on either a 1.5 T or 4 T scanner. Of note, the UCSF NIC scanner was updated during the study from a Siemens Magnetom Trio to a Magnetom Prisma model. To reduce the potential for confounding by this upgrade, images acquired on the Trio and Prisma models were treated statistically as coming from distinct scanners. Additional cohort details and the distribution of scans by scanner type are provided in Table 1.

All images were processed using FreeSurfer version 7.1 (Fischl et al., 2002; Desikan et al., 2006), manually checked for segmentation accuracy, and corrected as needed. We used FreeSurfer because it enables manual correction of segmentation errors in severely atrophied brains (Desikan et al., 2006); facilitates analysis of cortical thickness, which provides a more sensitive measure of atrophy than grey matter volume (Winkler et al., 2010); and enables the use of unique software packages not available on other imaging pipelines (Iglesias et al., 2018). Cortical regions of interest were defined using the Desikan-Killiany atlas (Desikan et al., 2006). Thalamic nuclei volumes were estimated using an extension of the FreeSurfer package (Iglesias et al., 2018), and all segmentations were manually checked. Cortical and thalamic illustrations of neuroimaging findings were generated using Freeview, the image-viewing software distributed with FreeSurfer.

### Experimental Design and Statistical Analysis

#### (i) General

Statistical analyses described below were completed using R version 4.1.2.

#### (ii) Whole thalamic volume analyses

Using multiple regression, we began our analyses by testing whether whole thalamic volumes were significantly different in *C9orf72* HRE carriers compared to normal controls, controlling for the effects of age, sex, CDR-SB score, education, MRI scanner type, and total intracranial volume (TIV). Following this, we tested whether total thalamic volumes varied by clinical diagnosis (normal control, pre-symptomatic *C9orf72* HRE carrier, MCI, FTD, or FTD-ALS) using the same covariates.

#### (iii) Thalamic nucleic volume analyses

Using FreeSurfer-estimated thalamic nuclei volumes, we next utilized hierarchical clustering to identify relationships between nucleic and clinical groupings. To ensure comparability across nuclei during hierarchical clustering, all nucleic volumes were normalized to TIV and z-scored prior to analysis. Hierarchical clustering analyses were performed, and the resulting heatmap was generated using the ComplexHeatmap package in R (Gu et al., 2016).

Based on the findings of the hierarchical clustering analyses, we next tested whether individual thalamic nucleic volumes significantly differed in *C9orf72* HRE carriers compared to normal controls, controlling for CDR-SB scores, age, sex, education, MRI scanner type, and TIV. As a sensitivity analysis to determine whether individual nuclei provided information independent of global thalamic atrophy, we next tested whether the nucleic volumes differed in HRE carriers versus controls, controlling for thalamic volumes, CDR-SB scores, age, sex, education, and MRI scanner type.

#### (iv) C9orf72 RNA expression analyses

We began the next stage of our analyses by confirming that *C9orf72* RNA expression was lower in peripheral blood samples from *C9orf72* HRE carriers relative to noncarriers, a finding suggested by prior literature. Multiple regression analysis was used to compare *C9orf72* RNA expression levels, covarying for age, sex, education, CDR-SB score, and batch.

To determine whether *C9orf72* expression provided clinically relevant, disease-related information, we tested whether it predicted CDR-SB scores, covarying for age, sex, education, and batch. Following this, we examined whether *C9orf72* expression provided information about clinical severity independent of thalamic atrophy in a combined multiple regression model, covarying for age, sex, education, MRI scanner type, TIV, and batch – using likelihood ratio testing to compare the combined model to models in which CDR-SB was predicted by *C9orf72* expression alone or thalamic volumes alone.

#### (v) Cortical thickness analyses

We concluded our neuroimaging analyses by examining associations between cortical thickness and four biomarkers of *C9orf72* HRE-related disease: clinical impairment as estimated by CDR-SB score, *C9orf72* HRE status, *C9orf72* expression, and volume of the top thalamic nucleus discovered in the above analyses. All cortical thicknesses were estimated using FreeSurfer as described above, and all analyses were performed using multiple regression models covarying for age, sex, education, MRI scanner type, and TIV. When analyzing *C9orf72* expression, batch was also included as a covariate.

### Peripheral Blood Mononuclear Cell (PBMC) Isolation

Blood samples from participants at UCSD diagnosed with ALS (both sporadic and due to *C9orf72* HRE) and healthy controls were collected into sodium heparin tubes and stored at room temperature for no longer than 30 hr from the time of draw. PBMCs were isolated via Ficoll density gradient centrifugation. Residual red blood cells were lysed in ammonium chloride-containing hemolytic buffer, then counted prior to freezing in 7% dimethyl sulfoxide in fetal bovine serum. PBMC samples were initially stored at -80°C, then transferred to a liquid nitrogen freezer within 72 hr.

### RNA Extraction

For whole-blood analyses, blood was drawn from participants at UCSF within 90 days of clinical assessment and stored in PAXgene blood RNA tubes (Qiagen) in liquid nitrogen. Briefly, total RNA was extracted from whole-blood samples using a MagMAX isolation kit (ThermoFisher) and RNA quality was assessed with a Bioanalyzer (Agilent). For PBMC samples, RNA was isolated via RNeasy kit (Qiagen). Samples with an RNA integrity score greater than 7 underwent library preparation for sequencing.

### RNA-seq

Library preparation and sequencing were performed at the UCLA Neuroscience Genomics core as previously described (Parikshak et al., 2016). The TruSeq Stranded Total RNA with Ribo-Zero Globin kit (Illumina) was used per the manufacturer’s protocol to prepare RNA for sequencing. Samples were sequenced in two batches: batch 1 (35 *C9orf72* HRE carriers, 10 sporadic ALS, 37 controls) was sequenced on a HiSeq 2500 generating 50 base pair paired-end reads, while batch 2 (24 *C9orf72* HRE carriers, 36 controls) was sequenced on a HiSeq 4000 generating 75 base pair paired-end reads. Samples in both batches were sequenced over multiple lanes and at an average depth of 50-60 M paired reads per sample.

### Sequencing Data Processing

Gene and transposable element (TE) abundance was determined in RNA-seq data as previously described (Jin and Hammell, 2018) using TEcount from the TEToolkit suite (Jin et al., 2015) (http://hammelllab.labsites.cshl.edu/software/). Reads were aligned to the GRCh38 build of the human reference genome using STAR v2.7.3a (Dobin et al., 2013) and a prebuilt GTF file of gene and TE annotation provided with TEtranscripts using the following parameters:

> STAR --runThreadN 8 --genomeDir /Index --readFilesIn sample_ID.R1.fastq.gz sample_ID.R2. fastq.gz --readFilesCommand zcat –outFileNamePrefix sample_ID --outSAMtype BAM Unsorted -- sjdbOverhang 100 --winAnchorMultimapNmax 200 --outFilterMultimapNmax 100 --sjdbGTFfile /Index/GRCh38_GENCODE_rmsk_TE.gtf

Gene and TE abundance were estimated from the resulting BAM files using TEcount, a comprehensive gene annotation GTF file from GENCODE release 34, and a prebuilt TE annotation index provided with TEtranscripts using the following parameters:

> TEcount -b sample_ID.Aligned.out.bam --format BAM --stranded reverse --mode multi --minL 1 -i 100 -- TE GRCh38_GENCODE_rmsk_TE.gtf --GTF gencode.v34.annotation.gtf –project sample_ID

### Differential Expression Analysis

Differential gene and TE expression was assessed using the DESeq2 package (Love et al., 2014) in R. Participant sex and age, and batch were included as covariates in the linear model when assessing differential expression of normalized counts in DESeq2. A change in gene or TE expression was considered statistically significant at a Benjamini-Hochberg false discovery rate (FDR)-adjusted *p* (*p*_FDR_) < 0.05.

### Gene Set Enrichment Analysis (GSEA)

Gene lists were generated from DESeq2 analyses and ranked by log_2_ fold-change (logFC). All pre-ranked lists were analyzed using the ‘GSEAPreranked’ tool in the GSEA software v4.1.0 (Mootha et al., 2003; Subramanian et al., 2005) together with curated, pre-generated gene sets in the Hallmark Molecular Signatures Database supplemented with gene sets representing the senescence-associated secretory and type I interferon (IFN-I) response pathways (De Cecco et al., 2019). Sample permutation (*n* = 10,000) was used to correct for multiple testing.

## Results

To explore the relationship between *C9orf72* HRE carrier status and thalamic atrophy, we began by assessing whole thalamic volumes according to gene carrier status and diagnosis. As expected, we confirmed results from prior studies (Sha et al., 2012; Bocchetta et al., 2018, 2020), finding that HRE carrier status was significantly associated with reduced total thalamic volume (Fig. 1A). When analyzed by diagnosis, presymptomatic *C9orf72* HRE carriers as well as those diagnosed with MCI and FTD showed significant reductions in total thalamic volume, while the smaller FTD-ALS group did not reach significance (Fig. 1B).

**Figure 1.**
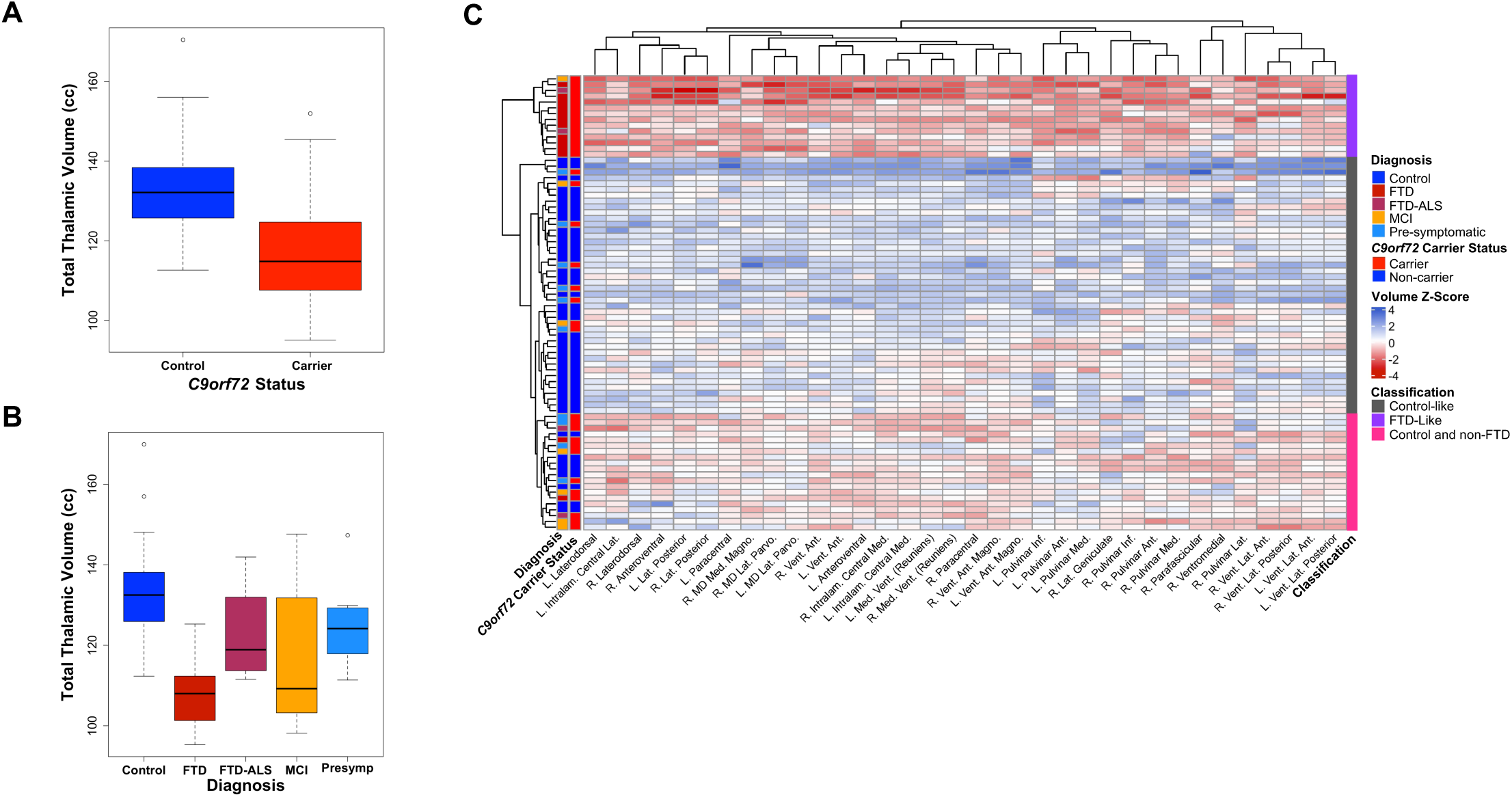
Comparisons of whole thalamic volume and thalamic nuclei volumes in *C9orf72* HRE carriers vs controls. ***A***, A box plot illustrates the results of a multiple regression analysis comparing total thalamic volume in *C9orf72* carriers of all diagnoses to controls. Across all diagnoses and accounting for the presence as well as severity of a disease (as measured by the Clinical Dementia Rating scale Sum of Boxes [CDR-SB] score), *C9orf72* carriers demonstrated lower total thalamic volumes compared to controls (*p* = 1.57×10^−6^). Additional covariates included in this analysis were age, sex, education, MRI scanner type (1.5 Tesla [T], 3 T, or 4 T), and TIV. ***B***, A box plot illustrates the results of a multiple regression analysis comparing total thalamic volumes in *C9orf72* HRE carriers vs controls by diagnosis. Of the four diagnostic groupings, three demonstrated a statistically significant difference vs controls: FTD (*p* = 0.005), MCI (*p* = 3.03×10^−6^), and pre-symptomatic (*p* = 0.003). The FTD-ALS group was not significantly different from controls (*p* = 0.16). Covariates included in this analysis were age, sex, education, MRI scanner type, CDR-SB score, and TIV. ***C***, Hierarchical clustering analyses of all thalamic nuclei volumes which significantly differed in *C9orf72* HRE carriers compared to controls at *p*_FDR_ < 0.05 are shown. The analysis revealed three primary groups: an FTD-like group (comprised almost exclusively of FTD cases, plus two FTD-ALS cases and an MCI case), a control-like group (comprised mostly of controls, plus pre-symptomatic *C9orf72* carriers, and an MCI case), and a more heterogeneous group consisting of controls and largely non-FTD cases (pre-symptomatic *C9orf72* carriers, MCI cases, and FTD-ALS cases). Abbreviations: Ant. – Anterior, Inf. – Inferior, Intralam. – Intralaminar, L. – Left, Lat. – Lateral, Magno. – Magnocellular, MD – Mediodorsal, Med. – Medial, Parvo. – Parvocellular, R. – Right, Vent. – Ventral. The covariates used in our multiple regression analyses comparing individual thalamic nuclei by *C9orf72* carrier status were the same as in (***B***).

To clarify the relationship between individual thalamic nuclei volumes and diagnosis, we performed hierarchical clustering following thalamic nuclei segmentation performed using an extension of the FreeSurfer pipeline (Iglesias et al., 2018). Hierarchical clustering revealed three broad groupings: an FTD-predominant group composed largely of HRE carriers diagnosed with FTD, a control-like group composed largely of non-carrier controls and several presymptomatic carriers, and a third, more heterogeneous group consisting largely of controls, presymptomatic carriers, and carriers with MCI (Fig. 1C). The FTD-predominant group showed a generally consistent atrophy pattern across 34 sub-thalamic regions that were significantly associated with *C9orf72* carrier status (Fig. 1C, Extended Data Table 1-1).

We next sought to determine whether any individual thalamic nuclei were disproportionately affected by *C9orf72* carrier status. We reasoned that the most atrophied thalamic nuclei in HRE carriers would remain significantly associated with *C9orf72* carrier status even after accounting for global thalamic atrophy. Intriguingly, we found that, after including total thalamic volume as a covariate in our regression analyses, the right mediodorsal lateral parvocellular (R MDl) nucleus showed highly significant atrophy (Fig. 2), consistent with a disproportionate effect on this nucleus. The R MDl nucleus projects to multiple regions of prefrontal cortex (PFC), dorsal anterior cingulate cortex (dACC), and frontal eye fields (FEF), and is thought to be involved in executive function as well as motor control of eye movements (Fig. 2, lower table) (Mitchell and Chakraborty, 2013; Ouhaz et al., 2018). Full results from the regression analyses described above are provided in Extended Data Tables 1-1 and 1-2.

**Figure 2.**
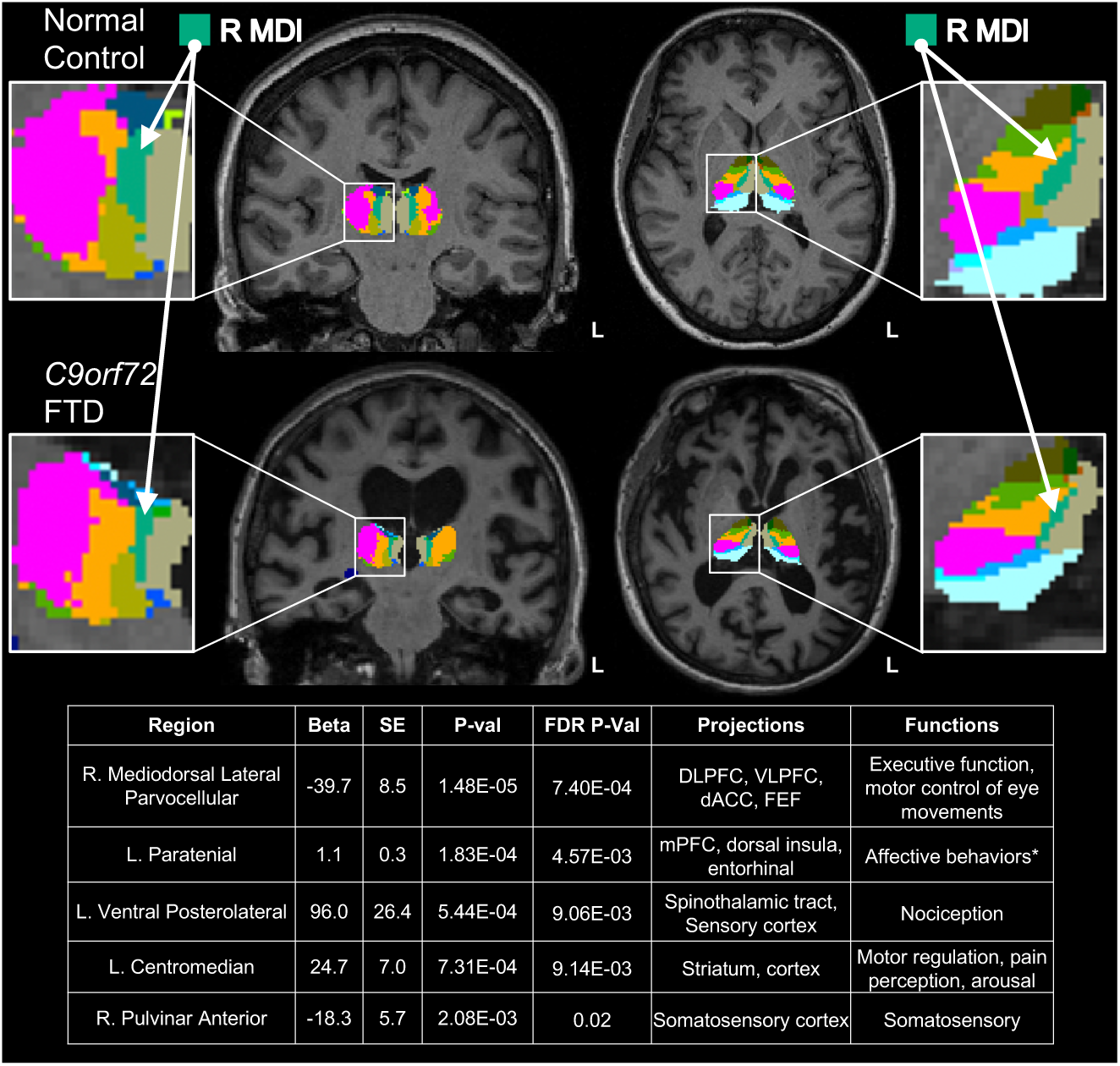
Mediodorsal lateral nucleus shows significant atrophy in *C9orf72* HRE carriers, independent of global thalamic atrophy. Five thalamic nuclei were significantly different in *C9orf72* HRE carriers vs controls at *p*_FDR_ < 0.05. Covariates in the multiple regression included total thalamic volume, age, sex, education, and MRI scanner type. The most significant association was observed in the right mediodorsal lateral parvocellular (R MDl) nucleus (*p* = 1.48 × 10^−5^). To illustrate the extent of volumetric differences observed in HRE carriers compared to controls, representative coronal and axial images (radiological orientation) from a normal healthy control (“Control”) and an FTD case (“*C9orf72* FTD”). The R MDl nucleus is shown in turquoise and indicated by white arrows in the insets. Of note, several nuclei that were significantly different in carriers vs controls, including the R MDl and left paratenial nuclei, are notable for connectivity to cortical regions frequently implicated in FTD (Vertes and Hoover, 2008; Mitchell and Chakraborty, 2013; Vertes et al., 2015; Ouhaz et al., 2018). Further, these nuclei are implicated in behavioral changes prominently impacted in C9*-*FTD, including executive function and affect (Mitchell and Chakraborty, 2013; Vertes et al., 2015; Ouhaz et al., 2018). Additional nuclei significantly associated with HRE carrier status include the left ventral posterolateral nucleus, which projects to the spinothalamic tract and sensory cortex and is involved in nociception (Al-Chaer et al., 1996; Darian-Smith et al., 1999; Krause et al., 2012); the left centromedian nucleus, which projects to motor cortex and is involved in motor regulation, pain perception, and arousal (Mai and Forutan, 2012; Ilyas et al., 2019); and the right pulvinar anterior nucleus, which projects to the somatosensory cortex and is involved in somatosensory function (Mai and Forutan, 2012).

Previous reports have demonstrated that peripheral blood expression of *C9orf72* is reduced in HRE carriers (Rizzu et al., 2016; McCauley et al., 2020). Using a novel bulk RNA-seq dataset generated from whole blood (described below), we confirmed a significant reduction in *C9orf72* RNA in HRE carriers in our neuroimaging cohort (Fig. 3A, Extended Data Table 1-3). Given that carriers showed both thalamic atrophy and reduced peripheral expression of *C9orf72*, we next asked whether thalamic volumes or peripheral *C9orf72* RNA levels were associated with clinical impairment, as measured by CDR-SB score. Strikingly, we found that both thalamic volumes and peripheral *C9orf72* levels were associated with CDR-SB scores (Fig. 3B). To determine whether these two factors independently predict clinical impairment, we included both variables in a combined regression model and found that both remained significant, indicating independent contributions. Likelihood ratio testing evaluating goodness-of-fit revealed that the combined model was superior to the models examining thalamic volumes alone (*p* = 0.04) or *C9orf72* expression alone (*p* = 2.14 × 10^−4^).

**Figure 3.**
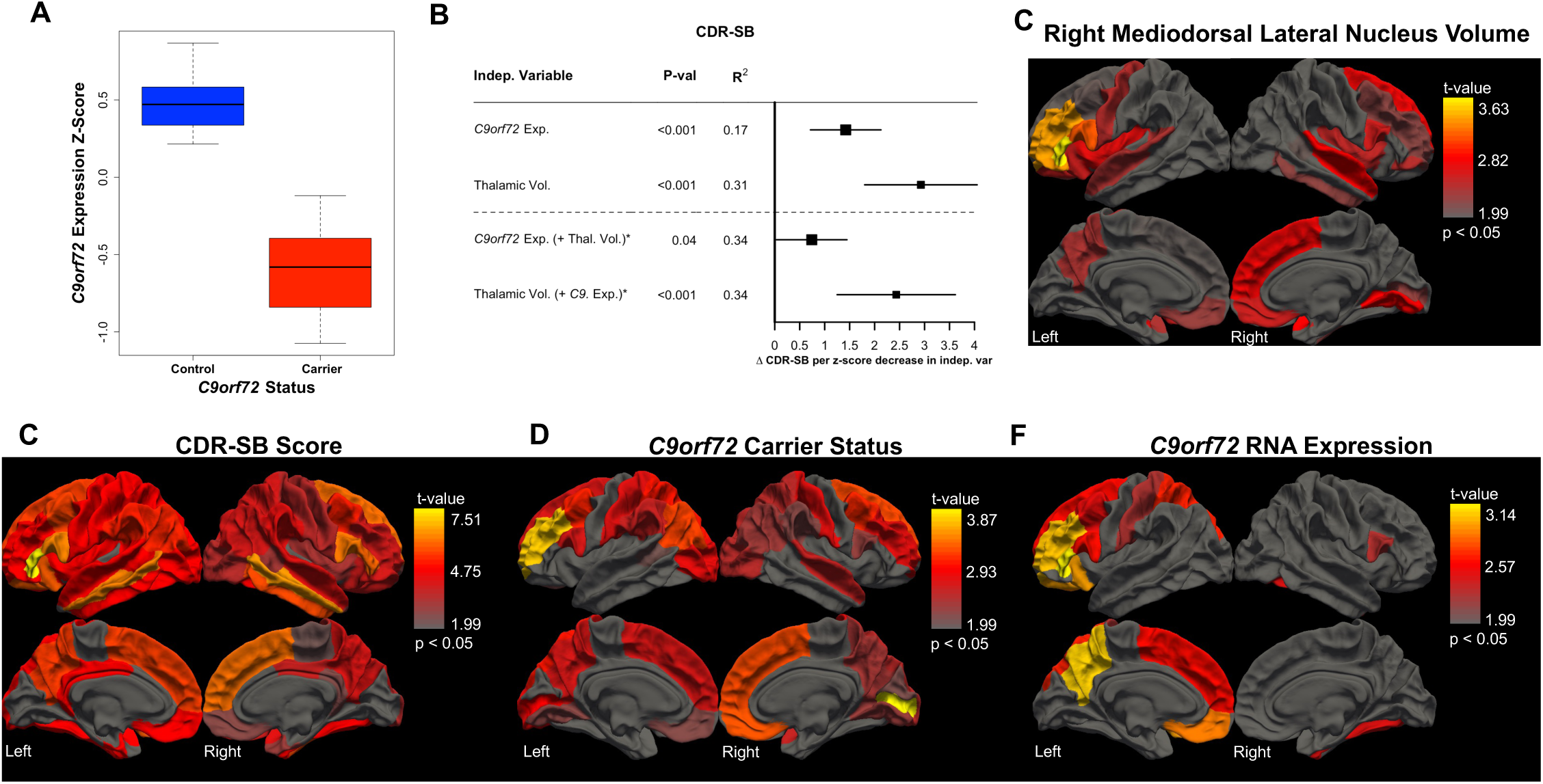
Peripheral *C9orf72* expression captures unique, disease-relevant information related to clinical severity. ***A***, *C9orf72* expression is significantly decreased in HRE carriers vs non-carrier controls (*p* = 5.56 × 10^−4^) in a multiple regression model covarying for age, sex, education, CDR-SB score, and sample processing batch. ***B***, *C9orf72* expression associates with clinical impairment, as measured by CDR-SB score (*p* = 1.84 × 10^−4^), even after correcting for age, sex, education, and batch. Given our finding of significantly lower total thalamic volumes in HRE carriers compared to controls (Fig. 1A; *p* = 1.57 × 10^−6^), we explored the relationship between *C9orf72* expression, thalamic volumes and CDR-SB scores. We found that thalamic volumes also predicted CDR-SB scores, again accounting for relevant covariates (*p* = 2.94 × 10^−6^; covariates: age, sex, education, MRI scanner type, and TIV). Both variables remained significant in a combined model: *C9orf72* expression (*p* = 0.04) and thalamic volumes (*p* = 1.47 × 10^−4^), suggesting that *C9orf72 expression* and thalamic volumes provide distinct, disease-relevant information. ***C***-***F***, Three-dimensional brain renderings depict results of multiple regression analyses (covarying for age, sex, education, clinical severity, MRI scanner type, and TIV) evaluating the relationship between cortical thickness and R MDl nucleus volumes (***C***), CDR-SB scores (***D***), *C9orf72* HRE carrier status (***E***), and *C9orf72* expression (***F***). ***C***, R MDl nucleus volumes associate with cortical thickness in multiple FTD-relevant regions such as prefrontal cortex and orbitofrontal cortex. ***D***, CDR-SB scores track global atrophy with relative sparing only of medial occipital cortex. ***E***, *C9orf72* carrier status associates with bifrontal thinning in HRE carriers with notable sparing of bilateral motor cortex. ***F***, *C9orf72* expression associates with left prefrontal cortex and left parietal cortex volumes with less prominent involvement of left orbitofrontal cortex and middle frontal gyrus.

We next asked whether R MDl nucleus volumes were associated with cortical thickness and found significant associations with multiple FTD-relevant regions, including PFC (Fig. 3C, Extended Data Table 1-4). As expected, we found that CDR-SB scores tracked global atrophy (Fig. 3D, Extended Data Table 1-5), while HRE carrier status was associated with bifrontal thinning with notable sparing of bilateral motor cortex (Fig. 3E, Extended Data Table 1-6). Finally, we found that peripheral *C9orf72* expression predicted left PFC and left parietal cortex volumes with less pronounced involvement of middle frontal gyrus (Fig. 3F, Extended Data Table 1-7).

Taken together, our neuroimaging analyses confirm global thalamic atrophy as well as significantly reduced peripheral *C9orf72* expression in HRE carriers. More significantly, we found that total thalamic volume and peripheral *C9orf72* expression levels independently predicted clinical impairment, and through parcellation of individual thalamic nuclei, we found that HRE carriers showed significant atrophy of the R MDl nucleus independent of global thalamic atrophy. Strikingly, we found that R MDl nucleus volumes, HRE carrier status, and peripheral *C9orf72* levels all associated with cortical thickness in FTD-relevant regions. The remarkable finding that peripheral *C9orf72* expression is associated with clinical impairment even after accounting for thalamic atrophy suggests that global peripheral transcriptomic changes in HRE carriers may capture additional disease-relevant biology. We therefore analyzed our whole-blood RNA-seq dataset with a particular focus on peripheral dysregulation of TEs due to their recent emergence in multiple neurodegenerative diseases along the FTD-ALS spectrum.

*C9orf72* HRE carriers (*n* = 49) and cognitively normal controls (*n* = 65) with available whole-blood-derived RNA were recruited for RNA-seq analyses (Table 2). HRE carriers had a range of clinical symptoms at the time of blood draw, with symptomatic participants having clinical diagnoses of MCI, FTD, ALS, and FTD-ALS. Also included were 12 presymptomatic HRE carriers who did not meet consensus criteria for any of the aforementioned clinical diagnoses. As expected, a significant difference in CDR-SB score by clinical diagnostic group was observed (*p* < 1×10^−5^). Presymptomatic HRE carriers were also significantly younger than both the control (*p* < 1×10^−5^) and symptomatic carriers (*p* < 1×10^−4^). There was no difference in sex across groups, and all participants self-reported race as “white.”

Total RNA derived from whole blood (*n* = 114) was deep sequenced to assess gene and TE expression across diagnostic groups. Since elevated TE expression—and in particular LINE1 element expression—has previously been reported in brain tissue from HRE carriers (Prudencio et al., 2017; Pereira et al., 2018; Tam et al., 2019; Zhang et al., 2019), we assessed total aggregate *L1HS* expression in cognitively normal older controls, and presymptomatic and symptomatic HRE carriers. After including sex, age, and batch as covariates in the differential expression (DE) analysis, we observed significantly elevated *L1HS* levels in whole blood from symptomatic carriers relative to controls (logFC = 1.1, *p*_FDR_ = 0.0001), while presymptomatic carriers had *L1HS* expression levels similar to controls (Fig. 4*C*). Beyond *L1HS*, nearly all detected TEs (1,109 out of 1,112) had significantly elevated expression in symptomatic HRE carriers, with the LTR subclass followed by the LINE superfamily making up the majority of the observed up-regulated TE expression (Fig. 4*A,B*). The latter observation is consistent with a general transcriptional de-repression of TEs, including the human-specific LINE-1 element, *L1HS*, detected in blood from symptomatic *C9orf72* HRE carriers.

**Figure 4.**
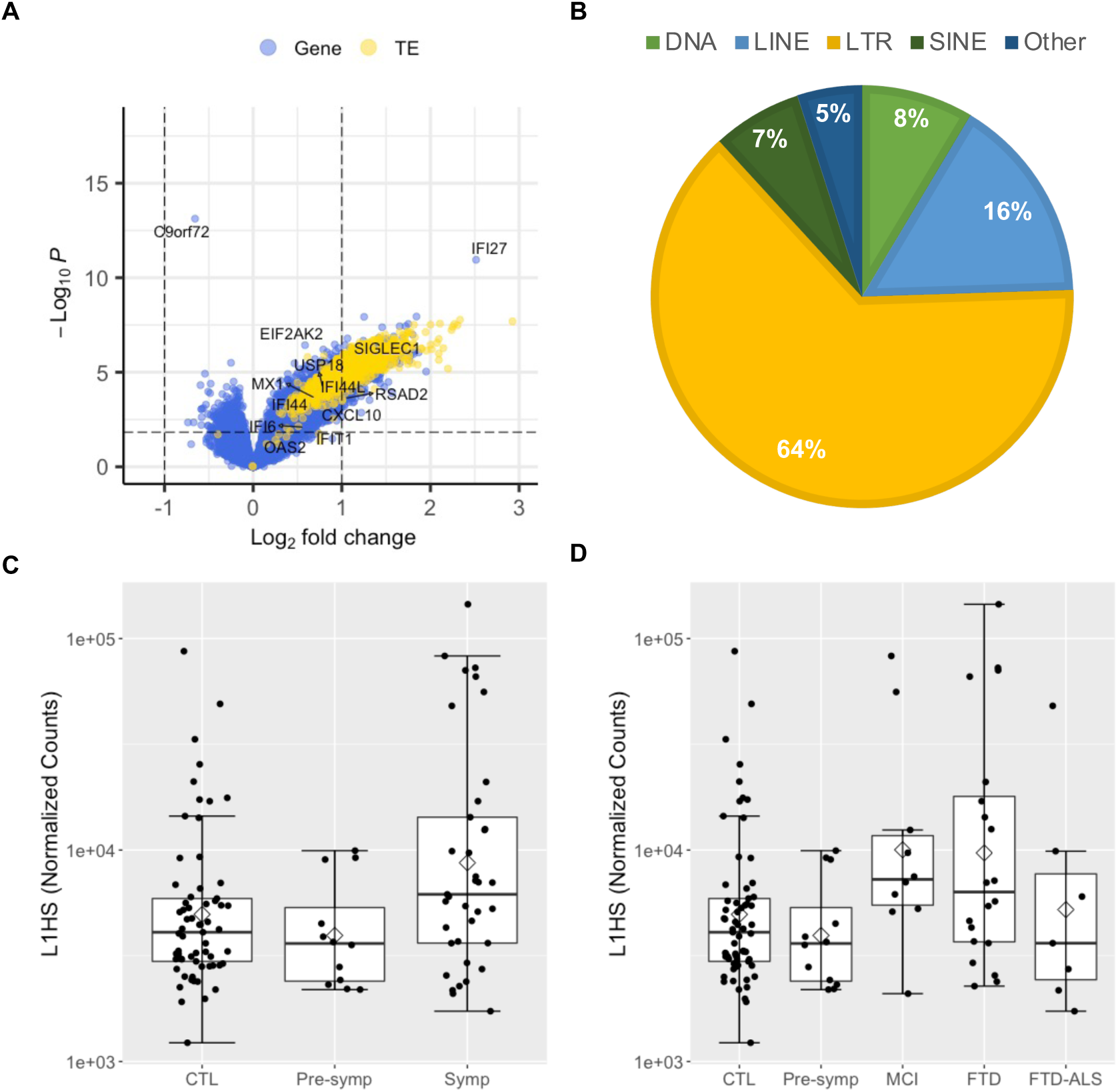
TE and type I interferon gene expression is elevated in symptomatic *C9orf72* HRE carrier whole blood. ***A***, Volcano plot of 17,220 genes 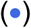 and 1,112 TEs 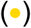 included in the differential expression (DE) analysis comparing symptomatic *C9orf72* HRE carriers (*n* = 37) to healthy controls (*n* = 65). Log_2_(fold-change) (logFC) and raw *p*-value determined by Wald test using DESeq2, including sex, age, and batch as covariates. Positive logFC indicates increased expression in *C9orf72* HRE carriers. The horizontal dashed line represents *p*_FDR_ = 0.05, with *p*_FDR_ < 0.05 considered statistically significant (i.e., data points above the horizontal dashed line, including *C9orf72*). Select type I interferon genes are highlighted. ***B***, the proportion of significantly elevated TEs in (***A***) by class and superfamily. ***C***, *L1HS* expression is significantly increased in symptomatic *C9orf72* HRE relative to control whole blood (logFC = 1.1, *p*_FDR_ = 0.0001). The mean (⋄) and median (—) are shown for each group. ***D***, *L1HS* expression by diagnostic group. *C9orf72* HRE carriers diagnosed with FTD and MCI have significantly elevated *L1HS* relative to controls (FTD: logFC = 1.22, *p*_FDR_ = 0.001; MCI: logFC = 1.30, *p*_FDR_ = 0.02). TE, transposable element; LINE, long interspersed nuclear element; LTR, long terminal repeat; SINE, short interspersed element; ALS, amyotrophic lateral sclerosis; CTL, healthy controls; FTD, frontotemporal dementia; MCI, mild cognitive impairment.

To determine whether specific diagnostic groups of symptomatic carriers underlie the observed increase in *L1HS* expression, we divided the symptomatic HRE carrier group by clinical diagnosis (Fig. 4*D*). Carriers diagnosed with an FTD-spectrum disorder or MCI had significantly elevated *L1HS* expression levels relative to cognitively normal older controls (FTD: logFC = 1.22, *p*_FDR_ = 0.001; MCI: logFC = 1.30, *p*_FDR_ = 0.02). No other clinical diagnostic group had statistically significant changes in *L1HS* expression relative to controls.

Changes in gene expression were also assessed in whole blood from HRE carriers. Out of 17,220 genes, expression of 731 and 3,606 genes was significantly decreased and increased, respectively, in symptomatic *C9orf72*+ participants (*p*_FDR_ < 0.05; Fig. 4*A*). Strikingly, *C9orf72* was the top differentially expressed gene (DEG) with reduced expression in symptomatic HRE carriers relative to controls (logFC = -0.66, *p*_FDR_ = 1.38×10^−09^; Fig. 4*A*). Presymptomatic carriers also had reduced *C9orf72* expression, but this difference did not reach statistical significance, likely due to the small sample size of this group (logFC = -0.45, *p*_raw_ = 0.005, *p*_FDR_ = 0.18). Since IFN-I signaling genes become activated by transcriptional de-repression of LINE-1 elements (De Cecco et al., 2019) and IFN-I signaling genes are upregulated in peripheral myeloid cells in *C9orf72* HRE carriers (McCauley et al., 2020), we further focused on these genes. Even though only a trend towards enrichment of the “Interferon Alpha Response” and “Type I Interferon Response” gene sets in symptomatic HRE carrier blood was observed (Extended Data Table 2-3), most genes in both pathways had increased expression (76 of 114 detected genes), including 27 with significantly increased expression and the second most significant DEG, *IFI27*, (*p*_FDR_ < 0.05; Fig. 4*A* and Extended Data Table 2-2). These findings support and extend previous observations of *C9orf72* HRE-associated patterns of gene expression dysregulation in peripheral blood cells by suggesting a possible role for *L1HS* in the observed transcriptional changes.

To determine if the above findings could be replicated in an independent cohort, we performed RNA-seq analyses on PBMCs isolated from *C9orf72* HRE carriers diagnosed with ALS (*n* = 10), individuals with sporadic ALS (*n* = 10), and healthy controls (*n* = 8). As expected, we again observed a significant reduction in *C9orf72* expression in HRE carriers compared to non-carriers (Fig. 5*A,B*). More importantly, we observed a global upregulation of TEs in C9-ALS compared to control PBMCs, a subset of which reached significance (Fig. 5*A*). In comparing C9-ALS to sporadic ALS, we observed a similar trend of increased TE expression, although most individual TEs did not reach significance (Fig. 5*B*). In particular, *L1HS* expression trended toward increased expression in C9-ALS relative to control PBMCs (logFC = 0.71, *p*_FDR_ = 0.075; Fig. 5*C*), suggesting de-repression of *L1HS* similar to that observed in whole-blood-derived RNA from symptomatic *C9orf72* HRE carriers (Fig. 4*C,D*). Finally, using GSEA, we identified significant enrichment of gene sets previously associated with *L1HS* activation (De Cecco et al., 2019), including the senescence-associated secretory pathway (SASP) and IFN-I response pathway (Fig. 5*D*).

**Figure 5.**
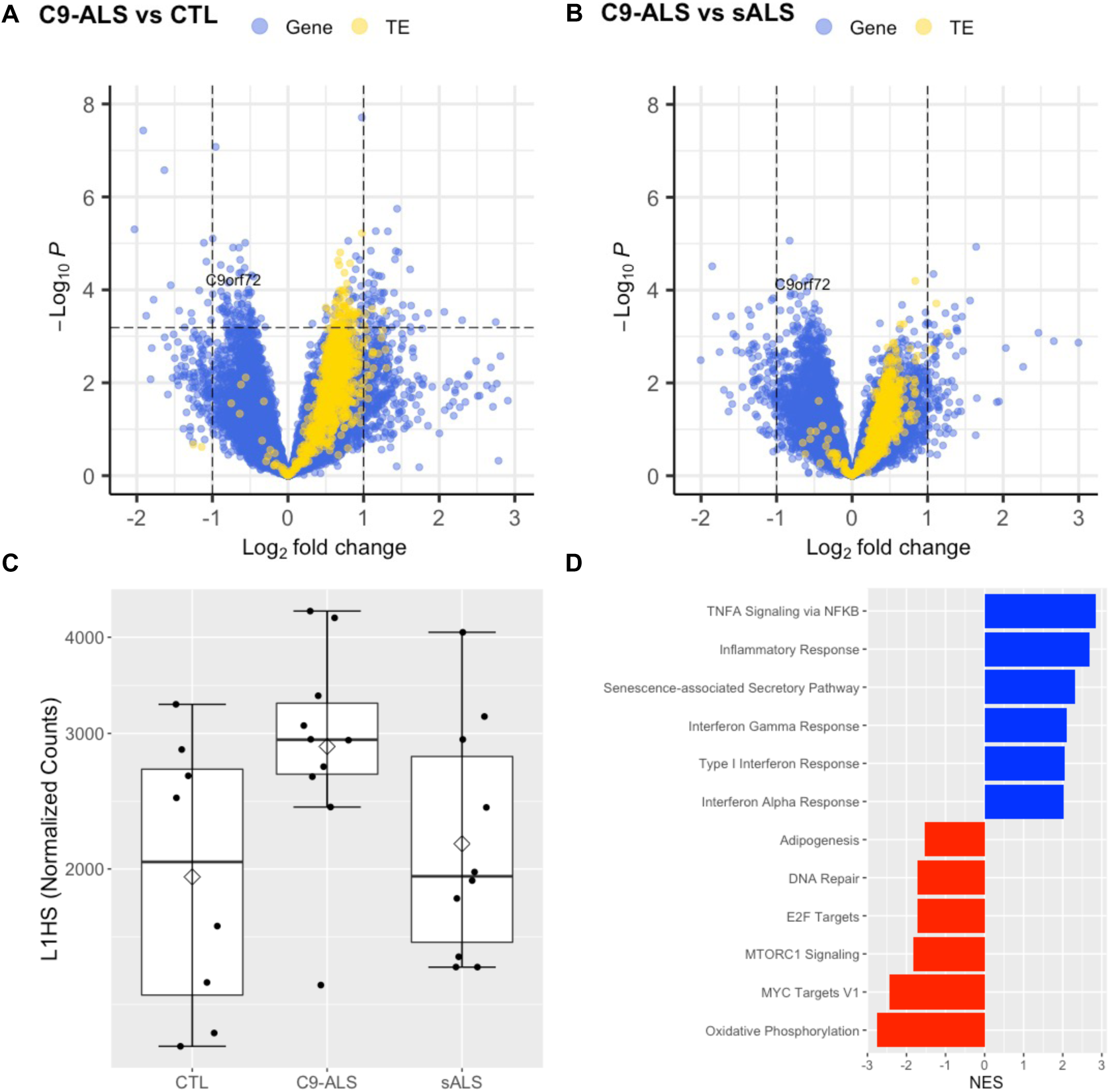
TE and type I interferon gene expression is elevated in PBMCs from *C9orf72* HRE carriers diagnosed with ALS. Volcano plot of 18,185 genes 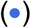 and 1,081 TEs 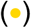 included in the DE analysis comparing *C9orf72*+ ALS (*n* = 10) to healthy control (*n* = 8) (***A***), or sporadic ALS (*n* = 10) (***B***) PBMCs. LogFC and raw *p*-values were determined by Wald test using DESeq2, including sex and age as covariates. Positive logFC indicates increased expression in *C9orf72* HRE carriers. The horizontal dashed line represents *p*_FDR_ = 0.05, with *p*_FDR_ < 0.05 considered statistically significant (i.e., data points above the horizontal dashed line, including *C9orf72*). ***C***, *L1HS* expression trends toward a significant increase in *C9orf72*+ ALS relative to control PBMCs (logFC = 0.71, *p*_FDR_ = 0.075). The mean (⋄) and median (—) are shown for each group. ***D***, GSEA results of the select enriched 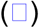 and suppressed 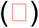 gene sets in *C9orf72*+ ALS relative to control PBMCs. Expression of genes in inflammatory pathways associated with *L1HS* activity are elevated in *C9orf72*+ ALS, including the interferon alpha response, type I interferon response, and senescence-associated secretory pathway gene sets. Full gene set enrichment results are in Extended Data Table 2-4. TE, transposable element; NES, normalized enrichment score.

Given the marked upregulation of *L1HS* observed in the peripheral blood of symptomatic HRE carriers, we finally asked whether peripheral *L1HS* levels could predict atrophy of thalamic nuclei in our neuroimaging cohort. Strikingly, we found that *L1HS* expression predicts volumes of several nuclei in the pulvinar (top three regions, *p*_raw_ < 7.9 × 10^−3^; Extended Data Table 1-8), a region of the thalamus previously shown to be affected in C9-FTD and implicated in motor function (Vatsavayai et al., 2016, 2018; Bocchetta et al., 2020).

## Discussion

Our radiogenomic integration of *C9orf72* HRE carrier neuroimaging and peripheral transcriptomic data enabled discoveries related to atrophy of specific thalamic nuclei and revealed an association between *C9orf72* expression and clinical impairment occurring independent of thalamic atrophy. In addition, we discovered globally up-regulated TE expression in peripheral blood of symptomatic HRE carriers in two independent cohorts. We also demonstrated strikingly increased expression of *L1HS* in affected HRE carriers and found that peripheral *L1HS* levels associated with thalamic nuclei volumes in FTD-relevant regions. Strikingly, our results indicate that de-repression of TE expression in C9-ALS/FTD patients is not restricted to the CNS. Moreover, the peripheral up-regulation of TEs such as *L1HS* we identified could be related to augmented expression of peripheral IFN-I signaling genes observed here and reported previously (De Cecco et al., 2019) (McCauley et al., 2020).

Our neuroimaging analyses confirmed previous findings of global thalamic atrophy in *C9orf72* HRE carriers (Sha et al., 2012; Bocchetta et al., 2018, 2020) and also identified atrophy of many individual thalamic nuclei, including those previously associated with HRE expansion (Lee et al., 2014, 2017; Yokoyama et al., 2014; Vatsavayai et al., 2016; Bocchetta et al., 2020). By controlling for total thalamic volume, we identified the R MDl nucleus as disproportionately atrophied in HRE carriers independent of global thalamic atrophy. The MDl nucleus projects to multiple regions of PFC, dACC, and FEF, and may influence executive function as well as motor control of eye movements (Mitchell and Chakraborty, 2013; Ouhaz et al., 2018). Given these connections, our group previously speculated that the MDl nucleus was a plausible candidate region for disproportionate dysfunction in C9-FTD (Yokoyama et al., 2014). In addition, presymptomatic *C9orf72* HRE carriers were recently shown to display dysfunctional saccadic eye movements (Behler et al., 2021), a function highly influenced by the FEF and superior colliculus (Purves and Williams, 2001). Our finding of greater MDl nucleus atrophy in HRE carriers suggests an intriguing possible mechanism connecting this emerging clinical biomarker of presymptomatic disease to a neuroanatomic region projecting to the FEF.

*C9orf72* HRE carriers and a subset of sporadic FTD/ALS cases show evidence of LINE1 element de-repression and retrotransposition, events thought to be related to loss of nuclear TDP-43 (Liu et al., 2019) (Prudencio et al., 2017) (Tam et al., 2019). In addition, heightened TE expression occurs in the context of tauopathies (Guo et al., 2018; Sun et al., 2018) and a distinct FTD-related proteinopathy due to pathogenic *CHMP2B* variation (Skibinski et al., 2005; Mackenzie and Neumann, 2016) (Fort-Aznar et al., 2020). Collectively, these findings suggest that TE de-repression may represent a general phenomenon occurring in multiple forms of neurodegeneration. However, to our knowledge, these findings are thus far reported exclusively within the CNS; we provide new evidence that TE up-regulation can also be measured in peripheral blood and may capture disease-relevant information.

Recent work in mice indicates a heightened inflammatory response in peripheral myeloid cells lacking *C9orf72* (McCauley et al., 2020). Consistent with these findings, *C9orf72* HRE in humans also appears to be associated with a heightened inflammatory response (Pinilla et al., 2021) driven by IFN-I signaling. Although the enhanced IFN-I signaling observed in C9-ALS myeloid cells is associated with a reduction in the peripheral expression of *C9orf72*, the primary mechanism driving inflammation is unknown. Impaired degradation of a promoter of IFN signaling has been proposed (McCauley et al., 2020), but gut microbiota (Burberry et al., 2020) and heightened expression of cytoplasmic double-stranded RNA (dsRNA) in C9-ALS/FTD brains may also be important contributors (Rodriguez et al., 2021). Our RNA-seq findings confirm the expected increase in IFN-I signaling gene expression in symptomatic HRE carriers (McCauley et al., 2020), and, importantly, also provide an additional plausible explanation for this phenomenon – the upstream transcriptional de-repression of TEs such as *L1HS*, which is known to trigger interferon signaling (Bourque et al., 2018).

Haploinsufficiency was suggested as a potential disease mechanism in C9-FTD/ALS in the initial reports characterizing *C9orf72* HRE (Dejesus-Hernandez et al., 2011; Renton et al., 2011; Gijselinck et al., 2012), and evidence in support of this idea soon emerged in model organisms (Ciura et al., 2013). Until recently, this mode of pathogenicity has received comparatively less attention than the proposed GOF mechanisms. However, the identification of C9orf72 as a key regulator of peripheral macrophage and brain microglia function in mice represented a key advance and suggested plausible mechanisms for haploinsufficiency (O’Rourke et al., 2016). Further, *in vitro* studies of induced motor neurons derived from C9-ALS patients suggest that haploinsufficiency may synergize with GOF mechanisms, ultimately leading to cell death (Shi et al., 2018). More recent work using mouse models supports the notion of pathogenic synergy between gains and loss of function (Zhu et al., 2020). Our findings demonstrating that *C9orf72* expression in peripheral blood captures information related to clinical severity, independent of an association with global thalamic atrophy, is potentially consistent with the notion that C9orf72 haploinsufficiency contributes to pathogenicity in the context of *C9orf72* HRE (Shi et al., 2018). We presume that peripheral blood levels of *C9orf72* relate to *C9orf72* expression in the CNS, although we have not directly tested this. Any potential pathologic effects of haploinsufficiency are likely to be mediated by haploinsufficient cells residing in the CNS, although a contributory role for haploinsufficient peripheral and/or infiltrative immune cells remains possible.

Building on these findings and hypotheses, we used a novel radiogenomic approach to identify specific thalamic nuclei atrophied in *C9orf72* HRE carriers, establish peripheral *C9orf72* expression as a potential predictor of clinical impairment independent of thalamic atrophy, and characterize the cortical correlates of R MDl atrophy. Prior work has demonstrated that the mediodorsal thalamus is atrophied in most genetic forms of FTD and that atrophy of the pulvinar is unique to *C9orf72* HRE carriers (Bocchetta et al., 2020), though these analyses were conducted covarying for TIV. Our analyses replicate and build on these findings by demonstrating that mediodorsal (i.e., R MDl nucleus) and pulvinar (i.e., R anterior pulvinar nucleus) atrophy remain significantly associated with *C9orf72* HRE status even after correcting for thalamic atrophy, a unique and important finding given the pronounced global thalamic atrophy observed in HRE carriers (Bocchetta et al., 2020; Spinelli et al., 2021). To our knowledge, our finding that peripheral *C9orf72* expression predicts clinical impairment independent of thalamic atrophy is novel and has potential not only as an adjunct to standard neuroimaging biomarkers of disease progression but also as a non-invasive biomarker that could be used to stratify patients in clinical trials or to measure treatment response. By characterizing the cortical regions most closely associated with R MDl atrophy, we identify atrophy in regions known to receive afferent connections from the MDl, possibly indicating degeneration of functionally connected regions.

Our study has several limitations. First, although we have replicated our primary RNA-seq findings – globally up-regulated peripheral TE expression and upregulation of IFN-I signaling genes – in symptomatic HRE carriers in an independent cohort, the replication cohort was smaller and therefore some findings replicated with only marginal significance (e.g., increased expression of *L1HS* in C9-ALS vs. control PBMCs). In addition, some of our neuroimaging findings were marginally significant (e.g., the association of peripheral *L1HS* levels with pulvinar nuclei volumes). It will therefore be important in future work not only to validate our RNA-seq findings but also to replicate our novel neuroimaging findings in large, independent, and diverse cohorts. Large-scale studies utilizing multimodal imaging (e.g., combined structural and functional MRI) will be required to precisely delineate the regions which atrophy symmetrically vs. asymmetrically and the underlying connectivity patterns that may drive our observations.

Our study raises many critical questions. First, why do TEs – and *L1HS* in particular – become de-repressed peripherally in symptomatic HRE carriers? This finding could be related to peripheral haploinsufficiency of C9orf72 protein, although CNS de-repression of TEs in sporadic neurodegenerative disease suggests an alternative mechanism. Loss of nuclear TDP-43 has been proposed as a potential mechanism, although this is unlikely to occur in the peripheral blood cells contributing to the signal in our RNA-seq data. Second, does peripheral *L1HS* up-regulation explain the upregulation of IFN-I signaling and SASP-related genes observed by us and others (De Cecco et al., 2019; McCauley et al., 2020)? Alternatively, is peripheral inflammation related more directly to crosstalk with CNS inflammatory processes (Bettcher et al., 2021)? Third, do elevated peripheral *L1HS* levels in symptomatic HRE carriers correlate with elevated levels of *L1HS* in the CNS? Fourth, can the peripheral elevation of *L1HS* or other TEs serve as a useful biomarker in *C9orf72* HRE carriers for prognosis or in clinical trials? Answers to these questions will be crucial for achieving a better understanding of *C9orf72* HRE-mediated disease and other forms of neurodegeneration.

## Supporting information

Extended Data - Tables 1-1 through 1-8

Extended Data - Tables 2-2 through 2-4

## Acknowledgments

Funding to JSY and for this project comes from NIH-NIA K01AG049152, R01AG062588, R01AG057234, P30AG062422, P01AG019724; NIH-NINDS U54NS123985; the AFTD Susan Marcus Memorial Fund; the Rainwater Charitable Foundation; the Larry L. Hillblom Foundation; the Bluefield Project to Cure Frontotemporal Dementia; the Alzheimer’s Association; the Global Brain Health Institute; the French Foundation; the Mary Oakley Foundation; and Transposon Therapeutics. SEL is supported by R01AG058233. VES is supported by R01AG052496. ALB receives research support from NIH (U19AG063911, R01AG038791, R01AG073482), the Tau Research Consortium, the Association for Frontotemporal Degeneration, Bluefield Project to Cure Frontotemporal Dementia, Corticobasal Degeneration Solutions, the Alzheimer’s Drug Discovery Foundation, and the Alzheimer’s Association, and has received research support from Biogen, Eisai and Regeneron.

We thank Giovanni Coppola for his longstanding support and collaboration.

## Notes

### Competing Interest Statement

ALB has served as a consultant for Aeovian, AGTC, Alector, Arkuda, Arvinas, AviadoBio, Boehringer Ingelheim, Denali, GSK, Life Edit, Humana, Oligomerix, Oscotec, Roche, Transposon, TrueBinding and Wave. JSY serves on the scientific advisory board for the Epstein Family Alzheimer Research Collaboration.

