## Extended Data - Tables 1-1 through 1-8 for "Radiogenomics of *C9orf72* expansion carriers reveals global transposable element de-repression and enables prediction of thalamic atrophy and clinical impairment"

Table 1-1: Thalamic volume differences in *C9orf72* HRE carriers compared to controls covarying for total intracranial volume

| Region | Beta | Standard Error | *P*-value | FDR *P*-Value |
| --- | --- | --- | --- | --- |
| R. Mediodorsal Lateral Parvocellular | -56.18 | 8.49 | 6.83E-09 | 3.42E-07 |
| R. Pulvinar Anterior | -38.21 | 6.78 | 3.61E-07 | 6.98E-06 |
| L. Ventral Anterior | -63.61 | 11.36 | 4.19E-07 | 6.98E-06 |
| R. Ventral Anterior | -58.40 | 11.91 | 6.12E-06 | 7.65E-05 |
| L. Lateral Posterior | -31.19 | 6.56 | 1.07E-05 | 1.07E-04 |
| L. Mediodorsal Lateral Parvocellular | -43.04 | 9.29 | 1.67E-05 | 1.31E-04 |
| R. Anteroventral | -33.59 | 7.29 | 1.83E-05 | 1.31E-04 |
| R. Intralaminar Central Medial | -13.59 | 2.99 | 2.33E-05 | 1.46E-04 |
| R. Ventral Anterior Magnocellular | -4.44 | 0.99 | 2.73E-05 | 1.52E-04 |
| L. Ventral Anterior Magnocellular | -4.26 | 0.96 | 3.41E-05 | 1.62E-04 |
| L. Intralaminar Central Medial | -13.52 | 3.05 | 3.56E-05 | 1.62E-04 |
| R. Ventral Lateral Anterior | -67.10 | 15.49 | 4.98E-05 | 1.92E-04 |
| L. Ventral Lateral Anterior | -69.35 | 16.01 | 4.99E-05 | 1.92E-04 |
| L. Anteroventral | -31.08 | 7.27 | 6.11E-05 | 2.18E-04 |
| R. Lateral Posterior | -27.71 | 6.55 | 7.23E-05 | 2.41E-04 |
| L. Laterodorsal | -10.53 | 2.73 | 2.52E-04 | 7.88E-04 |
| R. Pulvinar Medial | -144.58 | 37.70 | 2.77E-04 | 8.15E-04 |
| R. Pulvinar Inferior | -37.64 | 10.31 | 5.08E-04 | 1.34E-03 |
| R. Pulvinar Lateral | -33.73 | 9.24 | 5.08E-04 | 1.34E-03 |
| R. Mediodorsal Medial Magnocellular | -83.10 | 23.65 | 7.91E-04 | 1.98E-03 |
| L. Paracentral | -0.50 | 0.14 | 8.53E-04 | 2.03E-03 |
| L. Pulvinar Anterior | -23.45 | 6.83 | 1.02E-03 | 2.32E-03 |
| R. Ventral Lateral Posterior | -61.59 | 19.99 | 2.97E-03 | 6.46E-03 |
| R. Parafascicular | -6.55 | 2.15 | 3.28E-03 | 6.83E-03 |
| L. Medial Ventral (Reuniens) | -2.28 | 0.78 | 4.44E-03 | 8.88E-03 |
| R. Medial Ventral (Reuniens) | -2.63 | 0.91 | 5.06E-03 | 9.73E-03 |
| L. Pulvinar Inferior | -24.42 | 8.57 | 5.77E-03 | 0.01 |
| R. Lateral Geniculate | -24.84 | 9.01 | 7.47E-03 | 0.01 |
| L. Ventral Lateral Posterior | -57.51 | 21.21 | 8.46E-03 | 0.01 |
| R. Paracentral | -0.40 | 0.15 | 9.94E-03 | 0.02 |
| R. Ventromedial | -3.44 | 1.32 | 0.01 | 0.02 |
| L. Intralaminar Central Lateral | -6.16 | 2.40 | 0.01 | 0.02 |
| R. Laterodorsal | -8.09 | 3.21 | 0.01 | 0.02 |
| L. Pulvinar Medial | -81.61 | 34.24 | 0.02 | 0.03 |
| R. Intralaminar Central Lateral | -5.12 | 2.77 | 0.07 | 0.10 |
| L. Mediodorsal Medial Magnocellular | -50.21 | 28.64 | 0.08 | 0.11 |
| R. Ventral Posterolateral | -54.45 | 31.09 | 0.08 | 0.11 |
| L. Pulvinar Lateral | -16.61 | 9.62 | 0.09 | 0.12 |
| L. Suprageniculate | -3.49 | 2.11 | 0.10 | 0.13 |
| R. Suprageniculate | -2.76 | 1.86 | 0.14 | 0.17 |
| L. Lateral Geniculate | -13.54 | 9.21 | 0.15 | 0.17 |
| L. Parafascicular | -2.77 | 1.92 | 0.15 | 0.17 |
| R. Paratenial | -0.40 | 0.28 | 0.15 | 0.17 |
| R. Intralaminar Centromedian | -11.50 | 8.14 | 0.16 | 0.18 |
| L. Paratenial | 0.25 | 0.29 | 0.39 | 0.43 |
| L. Ventromedial | -0.87 | 1.16 | 0.45 | 0.49 |
| R. Medial Geniculate | -3.86 | 6.84 | 0.57 | 0.61 |
| L. Intralaminar Centromedian | -2.36 | 7.96 | 0.77 | 0.79 |
| L. Medial Geniculate | -1.55 | 5.32 | 0.77 | 0.79 |
| L. Ventral Posterolateral | -0.58 | 28.49 | 0.98 | 0.98 |

Comparisons of thalamic nuclei volumes in *C9orf72* HRE carriers vs. controls. Results from all 50 thalamic nuclei volumes estimated using Freesurfer 7.1 are shown above with *p*-values shown before and after FDR correction for multiple testing. All regression analysis covaried for clinical severity (as estimated by CDR-SB score), age, sex, education, MRI scanner type (1.5T, 3T, or 4T), and total intracranial volume. R. – Right, L. – Left.

Table 1-2: Thalamic volume differences in *C9orf72* HRE carriers compared to controls covarying for total thalamic volume

| Region | Beta | Standard Error | *P*-Value | FDR *P*-Value |
| --- | --- | --- | --- | --- |
| R. Mediodorsal Lateral Parvocellular | -39.72 | 8.51 | 1.48E-05 | 7.40E-04 |
| L. Paratenial | 1.12 | 0.28 | 1.83E-04 | 4.57E-03 |
| L. Ventral Posterolateral | 95.99 | 26.44 | 5.44E-04 | 9.06E-03 |
| L. Intralaminar Centromedian | 24.68 | 6.98 | 7.31E-04 | 9.14E-03 |
| R. Pulvinar Anterior | -18.30 | 5.72 | 2.08E-03 | 0.02 |
| R. Intralaminar Centromedian | 17.44 | 6.42 | 8.39E-03 | 0.07 |
| L. Ventral Anterior | -26.68 | 10.16 | 0.01 | 0.07 |
| R. Intralaminar Central Medial | -7.68 | 2.94 | 0.01 | 0.07 |
| L. Lateral Posterior | -16.26 | 6.46 | 0.01 | 0.07 |
| R. Anteroventral | -18.40 | 7.40 | 0.02 | 0.07 |
| L. Mediodorsal Lateral Parvocellular | -22.44 | 9.17 | 0.02 | 0.07 |
| L. Ventromedial | 2.64 | 1.09 | 0.02 | 0.07 |
| L. Intralaminar Central Medial | -7.26 | 3.04 | 0.02 | 0.08 |
| R. Ventral Posterolateral | 59.27 | 25.47 | 0.02 | 0.08 |
| L. Anteroventral | -16.13 | 7.29 | 0.03 | 0.10 |
| R. Ventral Anterior | -22.04 | 10.77 | 0.04 | 0.13 |
| L. Laterodorsal | -5.60 | 2.74 | 0.04 | 0.13 |
| R. Ventral Lateral Posterior | 27.00 | 13.44 | 0.05 | 0.13 |
| R. Pulvinar Lateral | -18.06 | 9.78 | 0.07 | 0.18 |
| L. Medial Geniculate | 9.35 | 5.37 | 0.09 | 0.20 |
| R. Lateral Posterior | -11.07 | 6.38 | 0.09 | 0.20 |
| L. Parafascicular | 3.14 | 1.82 | 0.09 | 0.20 |
| R. Paratenial | 0.42 | 0.25 | 0.10 | 0.21 |
| R. Ventral Anterior Magnocellular | -1.16 | 0.75 | 0.13 | 0.26 |
| R. Medial Ventral (Reuniens) | -1.43 | 0.95 | 0.14 | 0.27 |
| R. Medial Geniculate | 9.68 | 6.80 | 0.16 | 0.31 |
| L. Medial Ventral (Reuniens) | -1.11 | 0.80 | 0.17 | 0.31 |
| L. Pulvinar Medial | 38.05 | 27.97 | 0.18 | 0.32 |
| R. Pulvinar Inferior | -13.85 | 10.48 | 0.19 | 0.33 |
| L. Ventral Anterior Magnocellular | -1.01 | 0.79 | 0.20 | 0.34 |
| R. Mediodorsal Medial Magnocellular | -26.96 | 22.47 | 0.23 | 0.38 |
| L. Ventral Lateral Posterior | 19.03 | 16.91 | 0.26 | 0.41 |
| L. Ventral Lateral Anterior | -13.75 | 13.17 | 0.30 | 0.44 |
| L. Lateral Geniculate | 10.11 | 9.74 | 0.30 | 0.44 |
| R. Laterodorsal | -3.46 | 3.36 | 0.31 | 0.44 |
| L. Paracentral | -0.13 | 0.14 | 0.36 | 0.50 |
| L. Pulvinar Anterior | -5.59 | 6.21 | 0.37 | 0.50 |
| R. Pulvinar Medial | -23.80 | 32.71 | 0.47 | 0.62 |
| R. Ventromedial | 0.70 | 1.14 | 0.54 | 0.69 |
| L. Mediodorsal Medial Magnocellular | 13.89 | 27.99 | 0.62 | 0.78 |
| R. Paracentral | 0.05 | 0.13 | 0.69 | 0.84 |
| R. Ventral Lateral Anterior | -4.12 | 11.68 | 0.73 | 0.86 |
| L. Pulvinar Lateral | -3.09 | 10.20 | 0.76 | 0.86 |
| R. Suprageniculate | 0.56 | 1.90 | 0.77 | 0.86 |
| R. Intralaminar Central Lateral | 0.74 | 2.82 | 0.79 | 0.86 |
| L. Pulvinar Inferior | -2.31 | 8.77 | 0.79 | 0.86 |
| R. Parafascicular | -0.34 | 1.84 | 0.85 | 0.88 |
| L. Intralaminar Central Lateral | -0.41 | 2.38 | 0.86 | 0.88 |
| L. Suprageniculate | -0.33 | 2.20 | 0.88 | 0.88 |
| R. Lateral Geniculate | -1.38 | 9.45 | 0.88 | 0.88 |

Sensitivity analyses comparing of thalamic nuclei volumes in *C9orf72* HRE carriers vs. controls after covarying for total thalamic volume rather than total intracranial volume. Results from all 50 thalamic nuclei volumes estimated using Freesurfer 7.1 are shown above with *p*-values shown before and after FDR correction for multiple testing. All regression analysis covaried for clinical severity (as estimated by CDR-SB score), age, sex, education, MRI scanner type (1.5T, 3T, or 4T), and total thalamic volumes. R. – Right, L. – Left.

Table 1-3: *C9orf72* expression in *C9orf72* HRE carriers vs. controls

| Variable | Beta | Standard Error | *P*-Value |
| --- | --- | --- | --- |
| Sex (male) | 0.07 | 0.21 | 0.72 |
| Age (years) | -0.01 | 0.01 | 0.56 |
| Education (years) | -0.05 | 0.04 | 0.30 |
| CDR-SB score | -0.03 | 0.04 | 0.43 |
| Batch | -0.13 | 0.21 | 0.55 |
| *C9orf72* HRE status (carrier) | -1.09 | 0.30 | 5.56E-04 |

Multiple regression analyses demonstrate that *C9orf72* expression in *C9orf72* HRE carriers vs. controls is significantly decreased after covarying for the effects of sex, age, education, clinical severity (as estimated by CDR-SB score), and RNA-seq batch.

Table 1-4: Right mediodorsal lateral parvocellular nucleus volumes associate with cortical thicknesses

| Region | Beta | Standard Error | *P*-Value | FDR *P*-Value |
| --- | --- | --- | --- | --- |
| L. Pars Triangularis | 1.81E-03 | 4.99E-04 | 5.50E-04 | 0.03 |
| L. Rostral Middle Frontal | 1.70E-03 | 5.07E-04 | 1.30E-03 | 0.03 |
| L. Pars Orbitalis | 2.32E-03 | 7.00E-04 | 1.44E-03 | 0.03 |
| L. Pars Opercularis | 1.65E-03 | 5.30E-04 | 2.73E-03 | 0.05 |
| R. Superior Temporal | 1.31E-03 | 4.71E-04 | 6.79E-03 | 0.08 |
| R. Lingual | 1.14E-03 | 4.12E-04 | 7.06E-03 | 0.08 |
| R. Superior Frontal | 1.31E-03 | 4.79E-04 | 8.07E-03 | 0.08 |
| L. Insula | 1.53E-03 | 5.71E-04 | 9.25E-03 | 0.08 |
| R. Pars Opercularis | 1.28E-03 | 5.03E-04 | 0.01 | 0.08 |
| L. Lateral Orbitofrontal | 1.15E-03 | 4.56E-04 | 0.01 | 0.08 |
| R. Medial Orbitofrontal | 1.37E-03 | 5.43E-04 | 0.01 | 0.08 |
| R. Insula | 1.39E-03 | 5.62E-04 | 0.02 | 0.09 |
| L. Precentral | 1.31E-03 | 5.43E-04 | 0.02 | 0.09 |
| R. Pars Orbitalis | 1.67E-03 | 6.92E-04 | 0.02 | 0.09 |
| L. Superior Temporal | 1.24E-03 | 5.22E-04 | 0.02 | 0.09 |
| L. Precuneus | 1.11E-03 | 4.69E-04 | 0.02 | 0.09 |
| L. Medial Orbitofrontal | 1.17E-03 | 5.17E-04 | 0.03 | 0.10 |
| R. Pericalcarine | 1.23E-03 | 5.51E-04 | 0.03 | 0.10 |
| R. Inferior Temporal | 1.02E-03 | 4.59E-04 | 0.03 | 0.10 |
| R. Caudal Middle Frontal | 1.28E-03 | 5.82E-04 | 0.03 | 0.10 |
| R. Rostral Middle Frontal | 1.09E-03 | 4.96E-04 | 0.03 | 0.10 |
| L. Caudal Middle Frontal | 1.05E-03 | 5.12E-04 | 0.05 | 0.13 |
| R. Banks of the Superior Temporal Sulcus | 1.20E-03 | 5.91E-04 | 0.05 | 0.13 |
| L. Superior Frontal | 1.07E-03 | 5.30E-04 | 0.05 | 0.13 |
| L. Entorhinal | 2.13E-03 | 1.07E-03 | 0.05 | 0.13 |
| L. Postcentral | 9.24E-04 | 4.63E-04 | 0.05 | 0.13 |
| R. Paracentral | 1.05E-03 | 5.30E-04 | 0.05 | 0.13 |
| L. Lingual | 8.90E-04 | 4.57E-04 | 0.06 | 0.13 |
| R. Supramarginal | 1.01E-03 | 5.18E-04 | 0.06 | 0.13 |
| L. Banks of the Superior Temporal Sulcus | 8.90E-04 | 4.91E-04 | 0.07 | 0.17 |
| R. Temporal Pole | 2.31E-03 | 1.35E-03 | 0.09 | 0.20 |
| R. Precuneus | 7.68E-04 | 4.58E-04 | 0.10 | 0.20 |
| R. Middle Temporal | 8.69E-04 | 5.20E-04 | 0.10 | 0.20 |
| R. Postcentral | 8.22E-04 | 5.00E-04 | 0.10 | 0.21 |
| R. Lateral Orbitofrontal | 8.17E-04 | 5.01E-04 | 0.11 | 0.21 |
| R. Precentral | 1.02E-03 | 6.41E-04 | 0.12 | 0.22 |
| L. Temporal Pole | 2.07E-03 | 1.33E-03 | 0.12 | 0.23 |
| R. Pars Triangularis | 7.29E-04 | 4.76E-04 | 0.13 | 0.23 |
| R. Entorhinal | 1.93E-03 | 1.26E-03 | 0.13 | 0.23 |
| R. Cuneus | 7.18E-04 | 4.89E-04 | 0.15 | 0.25 |
| R. Inferior Parietal | 6.24E-04 | 4.57E-04 | 0.18 | 0.29 |
| L. Inferior Parietal | 4.90E-04 | 3.89E-04 | 0.21 | 0.34 |
| L. Posterior Cingulate | 7.16E-04 | 5.85E-04 | 0.23 | 0.36 |
| L. Lateral Occipital | 5.26E-04 | 4.45E-04 | 0.24 | 0.37 |
| R. Superior Parietal | 5.46E-04 | 4.72E-04 | 0.25 | 0.38 |
| L. Superior Parietal | 5.17E-04 | 4.50E-04 | 0.25 | 0.38 |
| L. Pericalcarine | 7.01E-04 | 6.26E-04 | 0.27 | 0.38 |
| L. Supramarginal | 5.52E-04 | 4.95E-04 | 0.27 | 0.38 |
| R. Fusiform | 4.14E-04 | 3.86E-04 | 0.29 | 0.39 |
| R. Transverse Temporal | 8.20E-04 | 7.69E-04 | 0.29 | 0.39 |
| R. Lateral Occipital | 4.33E-04 | 4.16E-04 | 0.30 | 0.40 |
| L. Cuneus | 5.99E-04 | 5.80E-04 | 0.31 | 0.40 |
| L. Fusiform | 3.72E-04 | 4.07E-04 | 0.36 | 0.47 |
| L. Middle Temporal | 4.05E-04 | 4.91E-04 | 0.41 | 0.52 |
| L. Parahippocampal | -5.41E-04 | 8.26E-04 | 0.51 | 0.64 |
| L. Rostral Anterior Cingulate | -3.29E-04 | 6.54E-04 | 0.62 | 0.75 |
| R. Isthmus Cingulate | -2.50E-04 | 5.64E-04 | 0.66 | 0.79 |
| L. Isthmus Cingulate | -2.61E-04 | 6.47E-04 | 0.69 | 0.79 |
| R. Posterior Cingulate | -1.87E-04 | 4.68E-04 | 0.69 | 0.79 |
| L. Inferior Temporal | 1.67E-04 | 4.28E-04 | 0.70 | 0.79 |
| R. Caudal Anterior Cingulate | -2.10E-04 | 7.08E-04 | 0.77 | 0.85 |
| R. Parahippocampal | -2.30E-04 | 8.16E-04 | 0.78 | 0.85 |
| L. Frontal Pole | 1.90E-04 | 8.29E-04 | 0.82 | 0.88 |
| R. Rostral Anterior Cingulate | -1.46E-04 | 7.38E-04 | 0.84 | 0.89 |
| L. Paracentral | -1.11E-04 | 5.99E-04 | 0.85 | 0.89 |
| R. Frontal Pole | 1.61E-04 | 9.96E-04 | 0.87 | 0.89 |
| L. Transverse Temporal | -1.20E-04 | 7.47E-04 | 0.87 | 0.89 |
| L. Caudal Anterior Cingulate | -3.15E-05 | 6.25E-04 | 0.96 | 0.96 |

Associations between right mediodorsal lateral parvocellular nucleus volumes and cortical thicknesses are shown. Results for all 68 cortical regions of interest from the Desikan-Killiany atlas with *p*-values shown before and after FDR correction for multiple testing. All regression analysis covaried for clinical severity (as estimated by CDR-SB score), age, sex, education, MRI scanner type (1.5T, 3T, or 4T), and total intracranial volume. R. – Right, L. – Left.

Table 1-5: CDR-SB score associations with whole-brain cortical thicknesses

| Region | Beta | Standard Error | *P*-Value | FDR *P*-Value |
| --- | --- | --- | --- | --- |
| L. Pars Triangularis | -3.55E-02 | 4.73E-03 | 1.57E-10 | 1.07E-08 |
| R. Middle Temporal | -2.99E-02 | 4.61E-03 | 1.12E-08 | 3.81E-07 |
| L. Pars Opercularis | -3.08E-02 | 4.92E-03 | 2.89E-08 | 5.34E-07 |
| L. Middle Temporal | -2.67E-02 | 4.29E-03 | 3.14E-08 | 5.34E-07 |
| R. Pars Opercularis | -2.80E-02 | 4.58E-03 | 5.09E-08 | 6.92E-07 |
| R. Superior Frontal | -2.63E-02 | 4.38E-03 | 7.85E-08 | 8.90E-07 |
| R. Pars Triangularis | -2.48E-02 | 4.20E-03 | 1.18E-07 | 1.15E-06 |
| L. Lateral Orbitofrontal | -2.40E-02 | 4.14E-03 | 1.79E-07 | 1.52E-06 |
| R. Inferior Temporal | -2.31E-02 | 4.13E-03 | 4.10E-07 | 3.10E-06 |
| L. Caudal Middle Frontal | -2.53E-02 | 4.58E-03 | 5.74E-07 | 3.90E-06 |
| L. Superior Frontal | -2.59E-02 | 4.74E-03 | 6.94E-07 | 4.29E-06 |
| L. Precuneus | -2.30E-02 | 4.24E-03 | 8.41E-07 | 4.77E-06 |
| L. Inferior Parietal | -1.77E-02 | 3.42E-03 | 2.22E-06 | 1.16E-05 |
| L. Supramarginal | -2.21E-02 | 4.34E-03 | 2.81E-06 | 1.31E-05 |
| R. Caudal Middle Frontal | -2.67E-02 | 5.24E-03 | 2.88E-06 | 1.31E-05 |
| R. Fusiform | -1.61E-02 | 3.38E-03 | 1.06E-05 | 4.51E-05 |
| L. Rostral Middle Frontal | -2.21E-02 | 4.76E-03 | 1.56E-05 | 6.24E-05 |
| L. Inferior Temporal | -1.71E-02 | 3.73E-03 | 1.85E-05 | 6.99E-05 |
| L. Pars Orbitalis | -3.00E-02 | 6.55E-03 | 2.05E-05 | 7.34E-05 |
| L. Superior Parietal | -1.78E-02 | 3.95E-03 | 2.69E-05 | 9.15E-05 |
| L. Superior Temporal | -2.10E-02 | 4.72E-03 | 3.26E-05 | 1.06E-04 |
| R. Supramarginal | -1.97E-02 | 4.62E-03 | 6.33E-05 | 1.96E-04 |
| L. Banks of the Superior Temporal Sulcus | -1.85E-02 | 4.37E-03 | 6.72E-05 | 1.97E-04 |
| R. Superior Temporal | -1.83E-02 | 4.32E-03 | 6.97E-05 | 1.97E-04 |
| L. Medial Orbitofrontal | -1.96E-02 | 4.66E-03 | 7.60E-05 | 2.07E-04 |
| R. Precuneus | -1.67E-02 | 4.06E-03 | 1.05E-04 | 2.75E-04 |
| L. Entorhinal | -3.89E-02 | 9.55E-03 | 1.24E-04 | 3.12E-04 |
| R. Superior Parietal | -1.67E-02 | 4.14E-03 | 1.37E-04 | 3.34E-04 |
| R. Inferior Parietal | -1.62E-02 | 4.02E-03 | 1.45E-04 | 3.40E-04 |
| L. Precentral | -1.96E-02 | 4.92E-03 | 1.61E-04 | 3.64E-04 |
| R. Pars Orbitalis | -2.49E-02 | 6.27E-03 | 1.68E-04 | 3.69E-04 |
| R. Rostral Middle Frontal | -1.77E-02 | 4.46E-03 | 1.76E-04 | 3.73E-04 |
| R. Lateral Orbitofrontal | -1.73E-02 | 4.44E-03 | 2.28E-04 | 4.69E-04 |
| L. Postcentral | -1.60E-02 | 4.14E-03 | 2.48E-04 | 4.96E-04 |
| R. Posterior Cingulate | -1.54E-02 | 4.07E-03 | 3.23E-04 | 6.28E-04 |
| R. Postcentral | -1.57E-02 | 4.43E-03 | 7.05E-04 | 1.33E-03 |
| R. Precentral | -1.98E-02 | 5.67E-03 | 8.27E-04 | 1.52E-03 |
| R. Lateral Occipital | -1.27E-02 | 3.64E-03 | 8.74E-04 | 1.56E-03 |
| R. Insula | -1.64E-02 | 5.09E-03 | 1.91E-03 | 3.33E-03 |
| R. Banks of the Superior Temporal Sulcus | -1.66E-02 | 5.28E-03 | 2.49E-03 | 4.23E-03 |
| R. Entorhinal | -3.31E-02 | 1.11E-02 | 4.07E-03 | 6.61E-03 |
| R. Temporal Pole | -3.55E-02 | 1.20E-02 | 4.08E-03 | 6.61E-03 |
| L. Insula | -1.44E-02 | 5.22E-03 | 7.44E-03 | 0.01 |
| R. Medial Orbitofrontal | -1.32E-02 | 4.93E-03 | 9.56E-03 | 0.01 |
| L. Posterior Cingulate | -1.37E-02 | 5.14E-03 | 9.71E-03 | 0.01 |
| R. Frontal Pole | -2.21E-02 | 8.66E-03 | 0.01 | 0.02 |
| R. Isthmus Cingulate | -1.26E-02 | 4.91E-03 | 0.01 | 0.02 |
| L. Lateral Occipital | -9.90E-03 | 3.91E-03 | 0.01 | 0.02 |
| L. Fusiform | -8.19E-03 | 3.55E-03 | 0.02 | 0.03 |
| L. Frontal Pole | -1.65E-02 | 7.20E-03 | 0.03 | 0.03 |
| R. Paracentral | -1.08E-02 | 4.74E-03 | 0.03 | 0.03 |
| L. Isthmus Cingulate | -1.25E-02 | 5.63E-03 | 0.03 | 0.04 |
| L. Paracentral | -9.84E-03 | 5.21E-03 | 0.06 | 0.08 |
| L. Temporal Pole | -2.02E-02 | 1.18E-02 | 0.09 | 0.11 |
| L. Cuneus | -8.27E-03 | 5.08E-03 | 0.11 | 0.13 |
| L. Lingual | -6.06E-03 | 4.08E-03 | 0.14 | 0.17 |
| R. Lingual | -5.50E-03 | 3.78E-03 | 0.15 | 0.18 |
| L. Pericalcarine | -6.59E-03 | 5.49E-03 | 0.23 | 0.27 |
| R. Cuneus | -5.07E-03 | 4.31E-03 | 0.24 | 0.28 |
| L. Parahippocampal | -7.29E-03 | 7.20E-03 | 0.31 | 0.36 |
| R. Rostral Anterior Cingulate | -5.73E-03 | 6.41E-03 | 0.37 | 0.42 |
| L. Rostral Anterior Cingulate | -4.21E-03 | 5.69E-03 | 0.46 | 0.51 |
| L. Caudal Anterior Cingulate | -3.69E-03 | 5.43E-03 | 0.50 | 0.54 |
| R. Parahippocampal | -4.69E-03 | 7.09E-03 | 0.51 | 0.54 |
| R. Transverse Temporal | -3.92E-03 | 6.73E-03 | 0.56 | 0.59 |
| R. Caudal Anterior Cingulate | -2.71E-03 | 6.15E-03 | 0.66 | 0.68 |
| L. Transverse Temporal | -1.68E-03 | 6.49E-03 | 0.80 | 0.81 |
| R. Pericalcarine | -8.15E-04 | 4.96E-03 | 0.87 | 0.87 |

Associations between CDR-SB score and cortical thicknesses are shown. Results for all 68 cortical regions of interest from the Desikan-Killiany atlas with associated *p*-values shown before and after FDR correction for multiple testing. All regression analysis covaried for age, sex, education, MRI scanner type (1.5T, 3T, or 4T), and total intracranial volume. R. – Right, L. – Left.

Table 1-6: Cortical thickness differences in *C9orf72* HRE carriers compared to controls

| Region | Beta | Standard Error | *P*-Value | FDR *P*-Value |
| --- | --- | --- | --- | --- |
| R. Pericalcarine | -0.18 | 0.05 | 2.45E-04 | 0.01 |
| L. Rostral Middle Frontal | -0.17 | 0.04 | 4.27E-04 | 0.01 |
| R. Medial Orbitofrontal | -0.15 | 0.05 | 1.76E-03 | 0.03 |
| R. Superior Frontal | -0.14 | 0.04 | 1.97E-03 | 0.03 |
| L. Inferior Parietal | -0.11 | 0.03 | 2.16E-03 | 0.03 |
| R. Caudal Middle Frontal | -0.16 | 0.05 | 2.94E-03 | 0.03 |
| L. Caudal Middle Frontal | -0.14 | 0.04 | 3.15E-03 | 0.03 |
| L. Superior Parietal | -0.12 | 0.04 | 3.62E-03 | 0.03 |
| L. Pericalcarine | -0.16 | 0.05 | 4.61E-03 | 0.03 |
| L. Lateral Occipital | -0.11 | 0.04 | 6.71E-03 | 0.04 |
| R. Postcentral | -0.12 | 0.04 | 7.17E-03 | 0.04 |
| R. Rostral Middle Frontal | -0.12 | 0.04 | 7.25E-03 | 0.04 |
| L. Postcentral | -0.11 | 0.04 | 7.60E-03 | 0.04 |
| L. Superior Frontal | -0.13 | 0.05 | 8.56E-03 | 0.04 |
| L. Precuneus | -0.11 | 0.04 | 8.80E-03 | 0.04 |
| L. Pars Opercularis | -0.13 | 0.05 | 8.97E-03 | 0.04 |
| L. Supramarginal | -0.11 | 0.04 | 0.01 | 0.04 |
| R. Superior Temporal | -0.11 | 0.04 | 0.01 | 0.04 |
| R. Superior Parietal | -0.10 | 0.04 | 0.01 | 0.05 |
| R. Lateral Occipital | -0.09 | 0.04 | 0.01 | 0.05 |
| R. Precuneus | -0.10 | 0.04 | 0.02 | 0.06 |
| R. Inferior Parietal | -0.09 | 0.04 | 0.02 | 0.07 |
| R. Lingual | -0.09 | 0.04 | 0.02 | 0.07 |
| R. Pars Opercularis | -0.10 | 0.05 | 0.02 | 0.07 |
| L. Lingual | -0.09 | 0.04 | 0.03 | 0.08 |
| R. Supramarginal | -0.10 | 0.05 | 0.03 | 0.08 |
| L. Medial Orbitofrontal | -0.10 | 0.05 | 0.03 | 0.08 |
| R. Cuneus | -0.09 | 0.04 | 0.03 | 0.08 |
| L. Banks of the Superior Temporal Sulcus | -0.09 | 0.04 | 0.04 | 0.09 |
| R. Paracentral | -0.10 | 0.05 | 0.05 | 0.11 |
| L. Posterior Cingulate | -0.10 | 0.05 | 0.05 | 0.11 |
| L. Lateral Orbitofrontal | -0.08 | 0.04 | 0.05 | 0.11 |
| L. Superior Temporal | -0.09 | 0.05 | 0.06 | 0.11 |
| L. Precentral | -0.10 | 0.05 | 0.06 | 0.11 |
| R. Inferior Temporal | -0.08 | 0.04 | 0.06 | 0.11 |
| L. Pars Triangularis | -0.09 | 0.05 | 0.08 | 0.15 |
| L. Pars Orbitalis | -0.11 | 0.07 | 0.11 | 0.20 |
| R. Middle Temporal | -0.07 | 0.05 | 0.12 | 0.20 |
| R. Transverse Temporal | -0.11 | 0.07 | 0.12 | 0.20 |
| R. Lateral Orbitofrontal | -0.07 | 0.05 | 0.12 | 0.20 |
| L. Insula | -0.08 | 0.05 | 0.13 | 0.21 |
| R. Pars Orbitalis | -0.10 | 0.06 | 0.13 | 0.21 |
| R. Temporal Pole | -0.18 | 0.12 | 0.14 | 0.22 |
| L. Entorhinal | -0.14 | 0.10 | 0.14 | 0.22 |
| L. Temporal Pole | -0.16 | 0.12 | 0.18 | 0.27 |
| L. Middle Temporal | -0.06 | 0.04 | 0.19 | 0.28 |
| R. Frontal Pole | -0.11 | 0.09 | 0.20 | 0.29 |
| R. Banks of the Superior Temporal Sulcus | -0.07 | 0.05 | 0.21 | 0.29 |
| R. Pars Triangularis | -0.05 | 0.04 | 0.21 | 0.29 |
| R. Precentral | -0.07 | 0.06 | 0.22 | 0.30 |
| R. Insula | -0.06 | 0.05 | 0.24 | 0.32 |
| L. Paracentral | -0.06 | 0.05 | 0.26 | 0.34 |
| L. Cuneus | -0.06 | 0.05 | 0.28 | 0.35 |
| L. Rostral Anterior Cingulate | -0.06 | 0.06 | 0.29 | 0.36 |
| R. Posterior Cingulate | -0.04 | 0.04 | 0.31 | 0.38 |
| L. Fusiform | -0.04 | 0.04 | 0.34 | 0.41 |
| R. Fusiform | -0.03 | 0.03 | 0.41 | 0.49 |
| L. Frontal Pole | -0.05 | 0.07 | 0.48 | 0.56 |
| R. Caudal Anterior Cingulate | 0.03 | 0.06 | 0.63 | 0.72 |
| L. Transverse Temporal | -0.03 | 0.07 | 0.66 | 0.74 |
| L. Parahippocampal | 0.02 | 0.07 | 0.82 | 0.90 |
| R. Entorhinal | -0.03 | 0.11 | 0.83 | 0.90 |
| R. Parahippocampal | -0.02 | 0.07 | 0.83 | 0.90 |
| L. Caudal Anterior Cingulate | 9.55E-03 | 0.06 | 0.87 | 0.92 |
| R. Isthmus Cingulate | -5.72E-03 | 0.05 | 0.91 | 0.95 |
| R. Rostral Anterior Cingulate | -6.01E-03 | 0.07 | 0.93 | 0.95 |
| L. Inferior Temporal | 2.97E-03 | 0.04 | 0.94 | 0.95 |
| L. Isthmus Cingulate | -3.76E-04 | 0.06 | 0.99 | 0.99 |

Associations between *C9orf72* HRE carrier status and cortical thicknesses are shown for all 68 cortical regions of interest from the Desikan-Killiany atlas with associated *p*-values shown before and after FDR correction for multiple testing. All regression analysis covaried for clinical severity (as estimated by CDR-SB score), age, sex, education, MRI scanner type (1.5T, 3T, or 4T), and total intracranial volume. R. – Right, L. – Left.

Table 1-7: Cortical thickness associations with *C9orf72* expression in HRE carriers and controls

| Region | Beta | Standard Error | *P*-Value | FDR *P*-Value |
| --- | --- | --- | --- | --- |
| L. Pars Triangularis | 0.10 | 0.03 | 2.54E-03 | 0.08 |
| L. Precuneus | 0.09 | 0.03 | 3.29E-03 | 0.08 |
| L. Rostral Middle Frontal | 0.09 | 0.03 | 3.61E-03 | 0.08 |
| L. Lateral Orbitofrontal | 0.08 | 0.03 | 4.48E-03 | 0.08 |
| L. Medial Orbitofrontal | 0.09 | 0.03 | 5.68E-03 | 0.08 |
| L. Superior Parietal | 0.07 | 0.03 | 0.01 | 0.10 |
| L. Pars Opercularis | 0.09 | 0.03 | 0.01 | 0.10 |
| L. Superior Frontal | 0.08 | 0.03 | 0.01 | 0.10 |
| L. Caudal Middle Frontal | 0.08 | 0.03 | 0.02 | 0.11 |
| R. Fusiform | 0.06 | 0.02 | 0.02 | 0.11 |
| R. Pars Opercularis | 0.07 | 0.03 | 0.03 | 0.15 |
| L. Postcentral | 0.06 | 0.03 | 0.03 | 0.15 |
| L. Pars Orbitalis | 0.09 | 0.05 | 0.06 | 0.26 |
| L. Frontal Pole | 0.10 | 0.05 | 0.06 | 0.26 |
| R. Rostral Middle Frontal | 0.06 | 0.03 | 0.06 | 0.26 |
| R. Superior Temporal | 0.06 | 0.03 | 0.06 | 0.26 |
| R. Middle Temporal | 0.06 | 0.03 | 0.07 | 0.26 |
| R. Pars Triangularis | 0.05 | 0.03 | 0.07 | 0.26 |
| R. Superior Parietal | 0.05 | 0.03 | 0.07 | 0.26 |
| L. Lateral Occipital | 0.05 | 0.03 | 0.08 | 0.29 |
| L. Pericalcarine | 0.06 | 0.04 | 0.09 | 0.29 |
| R. Superior Frontal | 0.05 | 0.03 | 0.10 | 0.29 |
| L. Supramarginal | 0.05 | 0.03 | 0.10 | 0.29 |
| R. Medial Orbitofrontal | 0.06 | 0.03 | 0.10 | 0.29 |
| R. Postcentral | 0.05 | 0.03 | 0.12 | 0.30 |
| R. Rostral Anterior Cingulate | -0.07 | 0.04 | 0.12 | 0.30 |
| L. Middle Temporal | 0.05 | 0.03 | 0.12 | 0.30 |
| R. Lingual | 0.04 | 0.03 | 0.13 | 0.32 |
| R. Inferior Parietal | 0.04 | 0.03 | 0.14 | 0.32 |
| R. Posterior Cingulate | 0.04 | 0.03 | 0.15 | 0.33 |
| L. Inferior Parietal | 0.03 | 0.02 | 0.17 | 0.37 |
| R. Caudal Middle Frontal | 0.05 | 0.04 | 0.18 | 0.37 |
| L. Superior Temporal | 0.04 | 0.03 | 0.19 | 0.37 |
| L. Precentral | 0.05 | 0.03 | 0.19 | 0.37 |
| L. Posterior Cingulate | 0.05 | 0.04 | 0.19 | 0.37 |
| L. Cuneus | 0.04 | 0.04 | 0.22 | 0.42 |
| R. Precuneus | 0.03 | 0.03 | 0.23 | 0.42 |
| R. Temporal Pole | 0.10 | 0.08 | 0.24 | 0.44 |
| L. Rostral Anterior Cingulate | 0.05 | 0.04 | 0.25 | 0.44 |
| R. Lateral Orbitofrontal | 0.04 | 0.03 | 0.26 | 0.44 |
| R. Banks of the Superior Temporal Sulcus | 0.04 | 0.04 | 0.27 | 0.44 |
| R. Entorhinal | 0.09 | 0.08 | 0.27 | 0.44 |
| L. Temporal Pole | 0.09 | 0.08 | 0.28 | 0.44 |
| L. Parahippocampal | -0.05 | 0.05 | 0.29 | 0.44 |
| R. Pericalcarine | 0.04 | 0.03 | 0.31 | 0.46 |
| L. Isthmus Cingulate | 0.04 | 0.04 | 0.31 | 0.46 |
| R. Lateral Occipital | 0.03 | 0.03 | 0.33 | 0.46 |
| R. Isthmus Cingulate | 0.03 | 0.03 | 0.33 | 0.46 |
| L. Banks of the Superior Temporal Sulcus | 0.03 | 0.03 | 0.37 | 0.51 |
| L. Fusiform | 0.02 | 0.03 | 0.42 | 0.55 |
| L. Inferior Temporal | 0.02 | 0.03 | 0.42 | 0.55 |
| L. Lingual | 0.02 | 0.03 | 0.43 | 0.55 |
| L. Insula | 0.03 | 0.04 | 0.43 | 0.55 |
| R. Pars Orbitalis | 0.03 | 0.04 | 0.44 | 0.55 |
| R. Paracentral | 0.02 | 0.03 | 0.49 | 0.61 |
| R. Inferior Temporal | 0.02 | 0.03 | 0.51 | 0.61 |
| L. Entorhinal | 0.04 | 0.07 | 0.53 | 0.64 |
| R. Supramarginal | 0.02 | 0.03 | 0.58 | 0.67 |
| L. Transverse Temporal | -0.03 | 0.05 | 0.59 | 0.67 |
| R. Insula | 0.02 | 0.04 | 0.59 | 0.67 |
| R. Caudal Anterior Cingulate | 0.02 | 0.04 | 0.63 | 0.71 |
| R. Parahippocampal | -0.02 | 0.05 | 0.73 | 0.80 |
| L. Paracentral | 0.01 | 0.04 | 0.78 | 0.84 |
| R. Transverse Temporal | 0.01 | 0.05 | 0.81 | 0.86 |
| R. Cuneus | 0.01 | 0.03 | 0.83 | 0.87 |
| L. Caudal Anterior Cingulate | 0.01 | 0.04 | 0.88 | 0.91 |
| R. Precentral | 4.39E-03 | 0.04 | 0.91 | 0.93 |
| R. Frontal Pole | -3.69E-03 | 0.06 | 0.95 | 0.95 |

Associations of *C9orf72* expression and cortical thicknesses are shown for all 68 cortical regions of interest from the Desikan-Killiany atlas with associated *p*-values shown before and after FDR correction for multiple testing. All regression analysis covaried for clinical severity (as estimated by CDR-SB score), age, sex, education, MRI scanner type (1.5T, 3T, or 4T), and total intracranial volume. R. – Right, L. – Left.

Table 1-8: Thalamic volume associations with *L1HS* expression

| Region | Beta | SE | *P*-Val | FDR *P*-val |
| --- | --- | --- | --- | --- |
| L. Pulvinar Lateral | -8.20 | 2.50 | 1.64E-03 | 0.08 |
| R. Pulvinar Medial | -30.30 | 10.95 | 7.31E-03 | 0.13 |
| R. Pulvinar Lateral | -7.08 | 2.58 | 7.86E-03 | 0.13 |
| R. Pulvinar Anterior | -5.60 | 2.18 | 0.01 | 0.16 |
| L. Pulvinar Medial | -21.77 | 9.51 | 0.03 | 0.25 |
| R. Lateral Posterior | -4.35 | 1.97 | 0.03 | 0.25 |
| L. Suprageniculate | -1.19 | 0.57 | 0.04 | 0.28 |
| L. Pulvinar Anterior | -4.02 | 2.00 | 0.05 | 0.30 |
| R. Ventromedial | -0.63 | 0.37 | 0.10 | 0.50 |
| L. Lateral Posterior | -3.38 | 2.04 | 0.10 | 0.50 |
| L. Ventromedial | -0.52 | 0.32 | 0.11 | 0.50 |
| L. Medial Geniculate | -2.23 | 1.45 | 0.13 | 0.52 |
| R. Pulvinar Inferior | -4.61 | 3.06 | 0.14 | 0.52 |
| R. Intralaminar Central Lateral | -1.10 | 0.77 | 0.16 | 0.55 |
| R. Medial Geniculate | -2.63 | 1.88 | 0.17 | 0.55 |
| L. Pulvinar Inferior | -3.27 | 2.45 | 0.19 | 0.57 |
| R. Suprageniculate | -0.68 | 0.52 | 0.19 | 0.57 |
| L. Ventral Lateral Anterior | -6.18 | 4.92 | 0.21 | 0.58 |
| R. Laterodorsal | -1.11 | 0.92 | 0.23 | 0.58 |
| R. Mediodorsal Lateral Parvocellular | -3.60 | 2.99 | 0.23 | 0.58 |
| L. Ventral Anterior | -4.44 | 3.76 | 0.24 | 0.58 |
| L. Ventral Anterior Magnocellular | -0.34 | 0.30 | 0.25 | 0.58 |
| L. Lateral Geniculate | -2.85 | 2.58 | 0.27 | 0.59 |
| R. Anteroventral | -2.37 | 2.20 | 0.28 | 0.59 |
| R. Ventral Anterior | -3.80 | 3.76 | 0.32 | 0.63 |
| L. Mediodorsal Lateral Parvocellular | -2.83 | 2.91 | 0.33 | 0.63 |
| R. Intralaminar Central Medial | -0.90 | 0.94 | 0.34 | 0.63 |
| L. Ventral Lateral Posterior | -5.44 | 6.12 | 0.38 | 0.63 |
| L. Parafascicular | 0.48 | 0.54 | 0.38 | 0.63 |
| L. Mediodorsal Medial Magnocellular | 6.92 | 7.91 | 0.38 | 0.63 |
| R. Lateral Geniculate | -2.25 | 2.60 | 0.39 | 0.63 |
| L. Laterodorsal | -0.67 | 0.83 | 0.42 | 0.66 |
| L. Medial Ventral (Reuniens) | -0.18 | 0.23 | 0.44 | 0.67 |
| L. Paratenial | 0.06 | 0.08 | 0.47 | 0.68 |
| R. Medial Ventral (Reuniens) | -0.19 | 0.27 | 0.48 | 0.68 |
| R. Ventral Posterolateral | -5.88 | 8.79 | 0.51 | 0.68 |
| L. Anteroventral | -1.48 | 2.24 | 0.51 | 0.68 |
| R. Ventral Anterior Magnocellular | -0.20 | 0.31 | 0.52 | 0.68 |
| R. Ventral Lateral Anterior | -2.86 | 4.81 | 0.55 | 0.71 |
| L. Ventral Posterolateral | -4.52 | 7.90 | 0.57 | 0.71 |
| R. Intralaminar Centromedian | -1.18 | 2.28 | 0.61 | 0.74 |
| L. Paracentral | -0.02 | 0.04 | 0.63 | 0.75 |
| R. Paracentral | 0.02 | 0.04 | 0.73 | 0.82 |
| R. Ventral Lateral Posterior | -2.07 | 5.91 | 0.73 | 0.82 |
| L. Intralaminar Central Medial | -0.32 | 0.96 | 0.74 | 0.82 |
| L. Intralaminar Centromedian | 0.68 | 2.21 | 0.76 | 0.83 |
| L. Intralaminar Central Lateral | -0.07 | 0.69 | 0.92 | 0.97 |
| R. Parafascicular | 0.05 | 0.63 | 0.94 | 0.97 |
| R. Mediodorsal Medial Magnocellular | -0.35 | 7.09 | 0.96 | 0.97 |
| R. Paratenial | 0.00 | 0.08 | 0.97 | 0.97 |

Associations of thalamic nuclei volumes with *L1HS* expression in a combined cohort of *C9orf72* HRE carriers and controls. Results from all 50 thalamic nuclei volumes estimated using Freesurfer 7.1 are shown above with *p*-values shown before and after FDR correction for multiple testing. All regression analysis covaried for clinical severity (as estimated by CDR-SB score), age, sex, education, MRI scanner type (1.5T, 3T, or 4T), and total intracranial volume. R. – Right, L. – Left.
