## Extended Data - Table 2-1 for "Radiogenomics of *C9orf72* expansion carriers reveals global transposable element de-repression and enables prediction of thalamic atrophy and clinical impairment"

**Table 2-1.** Demographic characteristics of PBMC RNA-seq cohort.

|  | Control | *C9orf72* HRE ALS | Sporadic ALS |
| --- | --- | --- | --- |
| *n* | 8 | 10 | 10 |
| Age, years (mean [SD]) | 51.5 (7.0) | 57.7 (8.4) | 51.3 (5.8) |
| Sex, *n* male (%) | 5 (62.5) | 4 (40.0) | 5 (50.0) |

ALS, amyotrophic lateral sclerosis. HRE, hexanucleotide repeat expansion.
